# LIFR recruits HGF-producing neutrophils to promote liver injury repair and regeneration

**DOI:** 10.1101/2023.03.18.533289

**Authors:** Yalan Deng, Zilong Zhao, Marisela Sheldon, Yang Zhao, Hongqi Teng, Consuelo Martinez, Jie Zhang, Chunru Lin, Yutong Sun, Fan Yao, Hao Zhu, Li Ma

## Abstract

The molecular links between tissue repair and tumorigenesis remain elusive. Here, we report that loss of the liver tumor suppressor *Lifr* in mouse hepatocytes impairs the recruitment and activity of reparative neutrophils, resulting in the inhibition of liver regeneration after partial hepatectomy or toxic injuries. On the other hand, overexpression of LIFR promotes liver repair and regeneration after injury. Interestingly, LIFR deficiency or overexpression does not affect hepatocyte proliferation *ex vivo* or *in vitro*. In response to physical or chemical damage to the liver, LIFR from hepatocytes promotes the secretion of the neutrophil chemoattractant CXCL1 (which binds CXCR2 to recruit neutrophils) and cholesterol in a STAT3-dependent manner. Cholesterol, in turn, acts on the recruited neutrophils to secrete hepatocyte growth factor (HGF) to accelerate hepatocyte proliferation and regeneration. Altogether, our findings reveal a LIFR-STAT3- CXCL1-CXCR2 axis and a LIFR-STAT3-cholesterol-HGF axis that mediate hepatic damage-induced crosstalk between hepatocytes and neutrophils to repair and regenerate the liver.

## Introduction

In response to chemical injury or surgical resection, the liver can regenerate to restore tissue homeostasis and organ function^1–3^. In chronic liver diseases, the regenerative and reparative capacity of the liver diminishes, and transplantation provides a curative option^4^. Due to the increasing incidence of fatty liver disease, cirrhosis, and liver cancer, interest in liver regeneration has been growing. Understanding the mechanisms of liver regeneration could be instrumental in preventing clinical complications associated with liver transplantation or resection for cancer removel^5, 6^. Moreover, since the number of liver donors is small, helping the liver to regenerate itself could offer an alternative treatment option for patients with advanced-stage liver disease.

Approximately 80% of the liver mass consists of hepatocytes, which are quiescent under normal physiological conditions. After chemical injuries or resection, hepatocytes re-enter the cell cycle. The induction of hepatocyte proliferation is regulated by various intracellular and extracellular factors^1, 2^ and is facilitated by the crosstalk between hepatocytes and hepatic immune cells^4, 7^. Among multiple signaling pathways involved, HGF-MET and EGFR (and its ligands) pathways are sufficient to induce hepatocyte proliferation directly^8–12^, and blockade of the MET or EGFR pathway abolishes liver regeneration after partial hepatectomy^13–17^. On the other hand, auxiliary mitogens, including interleukin (IL)-6 and tumor necrosis factor (TNF)-α, are insufficient to initiate hepatocyte proliferation on their own; rather, knockout of IL-6 or TNFR delays liver regeneration after partial hepatectomy^2, 18^.

After the liver injury, interactions between hepatocytes and immune cells contribute to liver repair and regeneration^4, 7^. Polymorphonuclear neutrophils are the largest population of circulating leukocytes and the first responder against invading pathogens^19, 20^. Accumulating evidence indicates that neutrophils can regulate liver injury repair and regeneration through several mechanisms^21^. For example, it has been shown that neutrophils promote liver regeneration by triggering Kupffer cell-dependent release of IL-6 and TNF-α^22^, or by inducing the phenotypic conversion of pro-inflammatory macrophages to reparative macrophages^23^. Although the contribution of neutrophils to tissue injury repair and regeneration is increasingly appreciated^23–27^, the mechanisms underlying neutrophil recruitment to the injured liver and the crosstalk between hepatocytes and neutrophils remain elusive.

Leukemia inhibitory factor receptor (LIFR) is expressed in various tissue types. Constitutive Lifr- knockout mice die within 24 hours of birth^28^, indicating that Lifr has an essential role in development. However, the functions of Lifr in many adult organs have not been described. *LIFR* expression is commonly downregulated in hepatocellular carcinoma (HCC)^29, 30^. Recently, we reported that hepatocyte-specific ablation of Lifr in mice (*Lifr*^fl/fl^;Alb-Cre) promoted hepatocarcinogen- and oncogene-induced HCC^29^. In the absence of hepatocarcinogen treatment or oncogene overexpression, *Lifr*^fl/fl^;Alb-Cre mice developed liver tumors by 2 years of age^29^.

Whether LIFR regulates liver tissue repair and regeneration is unknown. In this study, we found that loss of Lifr in hepatocytes impairs the recruitment and activity of reparative neutrophils, resulting in the inhibition of liver regeneration after injury. Mechanistically, in response to physical or chemical damage to the liver, hepatocytic LIFR promotes the secretion of the neutrophil chemoattractant CXCL1 and other pro-regenerative factors such as cholesterol in a STAT3- dependent manner. The recruited neutrophils, in turn, secrete the hepatocyte mitogen HGF to accelerate liver injury repair and regeneration.

## Results

### Loss of Lifr in hepatocytes impairs liver regeneration

After the removal of two-thirds of the liver mass in mice (a well-established method for studying liver regeneration^31^), Lifr protein levels fell initially but were restored at 48 hours, and after that, increased and then returned to basal levels at 2 weeks (**Extended Data Fig. 1a**). As expected, cyclin D1 and cyclin A2 were markedly upregulated during the recovery phase (**Extended Data Fig. 1a**). We suspected that the early downregulation of Lifr could be an initial response to acute damage, whereas the subsequent upregulation of Lifr may contribute to regeneration. To determine the function of Lifr in this process, we subjected liver-specific Lifr-knockout mice (*Lifr*^fl/fl^;Alb-Cre, as described in our recent publication^29^) and age-matched *Lifr*^fl/fl^ control animals to 2/3 partial hepatectomy (**Fig. 1a**). Relative to *Lifr*^fl/fl^ mice, *Lifr*^fl/fl^;Alb-Cre mice showed a lower liver-to-body weight ratio from 48 hours to 2 weeks after surgical resection (**Fig. 1b, c**), suggesting the compromised restoration of liver mass. Consistently, Lifr deficiency in the liver boosted hepatectomy-induced serum levels of alanine aminotransferase (ALT; **Fig. 1d**) and aspartate aminotransferase (AST; **Fig. 1e**), indicating increased liver injury. Moreover, *Lifr*^fl/fl^;Alb-Cre mice exhibited lower levels of hepatocyte proliferation than *Lifr*^fl/fl^ mice after hepatectomy, as evidenced by fewer mitotic nuclei (**Fig. 1f, g**) and Ki67-positive hepatocytes (**Fig. 1h, i**) as well as lower cyclin D1, cyclin A2, cyclin B1, and cyclin E1 levels (**Extended Data Fig. 1b, c**).

**Figure 1.**
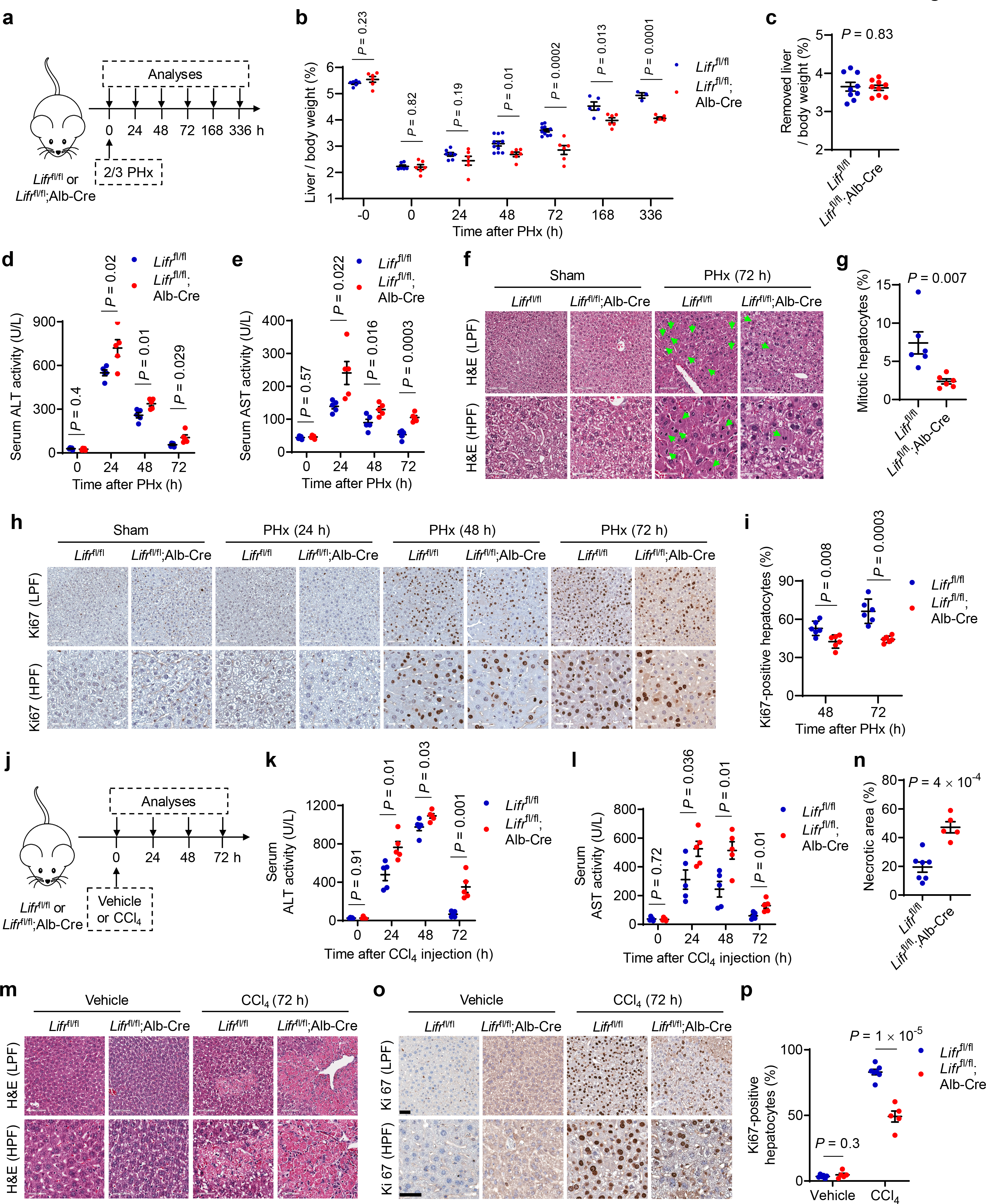
Loss of Lifr in hepatocytes impairs liver regeneration. **a.** Schematic of the experimental design for panels **b-i**. **b.** Liver-to-body weight ratio of *Lifr*^fl/fl^ and *Lifr*^fl/fl^;Alb-Cre mice at the indicated time points after 2/3 partial hepatectomy (PHx). *n* = 6, 6, 6, 6, 7, 6, 11, 6, 10, 6, 5, 6, 3, and 5 mice. **c.** Ratio of removed liver weight to body weight. *n* = 9 mice per group. **d**, **e.** Serum ALT (**d**) and AST (**e**) levels in *Lifr*^fl/fl^ and *Lifr*^fl/fl^;Alb-Cre mice at the indicated time points after PHx. *n* = 5 mice per group. **f**, **g.** Representative H&E staining (**f**) and percentage of mitotic hepatocytes (**g**; green arrows indicate mitotic nuclei) in the livers of *Lifr*^fl/fl^ and *Lifr*^fl/fl^;Alb-Cre mice at 72 hours after PHx. LPF: low-power field; HPF: high-power field. Scale bars, 100 μm in upper panels and 50 μm in lower panels. *n* = 6 mice per group. **h**, **i.** Immunohistochemical staining of Ki67 (**h**) and percentage of Ki67-positive hepatocytes (**i**) in the livers of *Lifr*^fl/fl^ and *Lifr*^fl/fl^;Alb-Cre mice at the indicated times after PHx. LPF: low-power field; HPF: high-power field. Scale bars, 100 μm in upper panels and 50 μm in lower panels. *n* = 6 mice per group. **j.** Schematic of the experimental design for panels **k-p**. **k**, **l.** Serum ALT (**k**) and AST (**l**) levels in *Lifr*^fl/fl^ and *Lifr*^fl/fl^;Alb-Cre mice at the indicated times after CCl4 treatment. *n* = 5 mice per group. **m**, **n.** Representative H&E staining (**m**) and percentage of necrotic areas (**n**) in the livers of *Lifr*^fl/fl^ and *Lifr*^fl/fl^;Alb-Cre mice at 72 hours after CCl4 treatment. LPF: low-power field; HPF: high-power field. Scale bars, 100 μm in upper panels and 50 μm in lower panels. *n* = 7 and 5 mice. **o**, **p.** Immunohistochemical staining of Ki67 (**o**) and percentage of Ki67-positive hepatocytes (**p**) in the livers of *Lifr*^fl/fl^ and *Lifr*^fl/fl^;Alb-Cre mice at 72 hours after CCl4 treatment. LPF: low-power field; HPF: high-power field. Scale bars, 50 μm. *n* = 6, 5, 7, and 5 mice. Statistical significance in **b-e**, **g**, **i**, **k**, **l**, **n**, and **p** was determined by a two-tailed unpaired *t*-test. Error bars are s.e.m.

To further test Lifr loss on liver repair and regeneration, we performed intraperitoneal injection of carbon tetrachloride (CCl4), which causes centrilobular hepatocyte injury^32, 33^ (**Fig. 1j**). Compared with *Lifr*^fl/fl^ mice, *Lifr*^fl/fl^;Alb-Cre mice showed impaired liver injury repair after CCl4 treatment, as gauged by serum levels of ALT and AST (**Fig. 1k, l**), necrotic areas (**Fig. 1m, n**), and TUNEL staining (**Extended Data Fig. 1d, e**). Moreover, *Lifr*^fl/fl^;Alb-Cre mice exhibited less hepatocyte proliferation, as evidenced by fewer Ki67-positive hepatocytes (**Fig. 1o, p**). qPCR analysis of the liver tissues of CCl4-treated *Lifr*^fl/fl^ and *Lifr*^fl/fl^;Alb-Cre mice showed that Lifr loss in hepatocytes impeded injury-induced upregulation of cyclin D1, cyclin A2, and proliferating cell nuclear antigen (PCNA) (**Extended Data Fig. 1f**). Taken together, these findings demonstrate that hepatocytic Lifr deficiency delays liver injury repair and regeneration after surgical resection or chemical damage of the mouse liver.

### Overexpression of LIFR promotes liver injury repair and regeneration

To determine whether Lifr overexpression promotes liver injury repair and regeneration, we injected C57BL/6J mice via the tail vein with control adenovirus or LIFR-expressing adenovirus^29^ 5 days before treatment with vehicle or CCl4, and conducted analyses 48 hours after the treatment (**Fig. 2a**). Adenoviral LIFR delivery substantially accelerated liver injury repair after CCl4 treatment, as gauged by serum levels of ALT and AST (**Fig. 2b, c**), necrotic areas (**Fig. 2d, e**), and TUNEL staining (**Fig. 2f, g**). Moreover, LIFR overexpression enhanced hepatocyte proliferation after CCl4-induced injury, as evidenced by a significant increase in the percentage of Ki67-positive hepatocytes (**Fig. 2h, i**).

**Figure 2.**
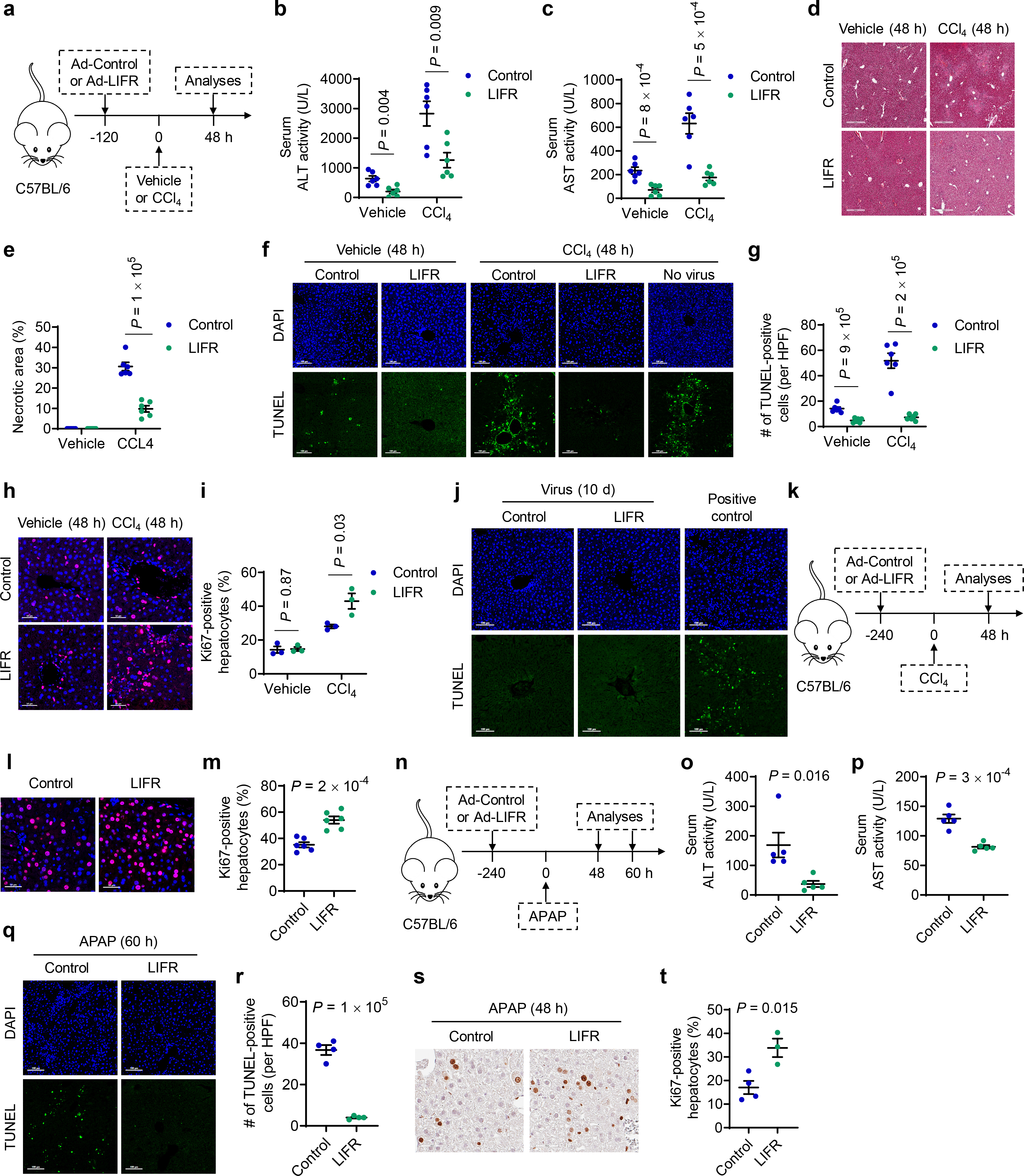
Overexpression of LIFR promotes liver injury repair and regeneration. **a**-**i.** C57BL/6J mice received control or LIFR-expressing adenovirus 5 days before CCl4 or vehicle treatment. Analyses were done at 48 hours after treatment. **a.** Schematic of the experimental design. **b**, **c.** Serum ALT (**b**) and AST (**c**) levels in mice after CCl4 or vehicle treatment. *n* = 6 mice per group. **d**, **e.** Representative H&E staining (**d**) and percentage of necrotic areas (**e**) in mouse livers after CCl4 or vehicle treatment. Scale bars, 500 μm. *n* = 6 mice per group. **f**, **g.** Representative DAPI and TUNEL staining (**f**) and the number of TUNEL-positive hepatocytes per high-power field (HPF; **g**) in mouse livers after CCl4 or vehicle treatment. Scale bars, 100 μm. *n* = 6 mice per group. **h**, **i.** Immunofluorescence staining of Ki67 (**h**; overlay with DAPI staining) and percentage of Ki67-positive hepatocytes (**i**) in mouse livers after CCl4 or vehicle treatment. Scale bars, 50 μm. *n* = 3 mice per group. **j.** Representative DAPI and TUNEL staining of mouse livers 10 days after injection of control or LIFR-expressing adenovirus. Scale bars, 100 μm. **k**-**m.** C57BL/6J mice received control or LIFR-expressing adenovirus 10 days before CCl4 treatment. Analyses were done at 48 hours after treatment. **k.** Schematic of the experimental design. **l**, **m.** Immunofluorescence staining of Ki67 (**l**; overlay with DAPI staining) and percentage of Ki67-positive hepatocytes (**m**) after CCl4 treatment. Scale bars, 50 μm. *n* = 6 mice per group. **n**-**t.** C57BL/6J mice received control or LIFR-expressing adenovirus 10 days before acetaminophen (APAP) treatment. Analyses were done at 48 or 60 hours after APAP treatment. **n.** Schematic of the experimental design. **o**, **p.** Serum ALT (**o**) and AST (**p**) levels in mice after APAP treatment. *n* = 5 mice per group. **q**, **r.** Representative TUNEL staining (**q**) and the number of TUNEL-positive hepatocytes per high-power field (HPF; **r**) in mouse livers after APAP treatment. Scale bars, 100 μm. *n* = 4 mice per group. **s**, **t.** Immunohistochemical staining of Ki67 (**s**) and percentage of Ki67-positive hepatocytes (**t**) after APAP treatment. Scale bars, 50 μm. *n* = 4 and 3 mice. Statistical significance in **b**, **c**, **e**, **g**, **i**, **m**, **o**, **p**, **r**, and **t** was determined by a two-tailed unpaired *t*- test. Error bars are s.e.m.

It should be noted that without CCl4 treatment, the injection of adenovirus itself caused mild liver damage, which was alleviated by LIFR overexpression (**Fig. 2b, c, f, g**; the vehicle group). To test the effect of adenoviral LIFR delivery in a condition that separates the repair of CCl4-induced liver damage from the repair of adenovirus-induced injury, we injected mice via the tail vein with control or LIFR-expressing adenovirus 10 days before CCl4 treatment, based on our observation that virus-induced liver damage disappeared at 10 days after adenovirus injection (**Fig. 2j**). Consistently, Ki67 staining results indicated that overexpression of LIFR in this setting boosted hepatocyte proliferation after CCl4-induced liver damage (**Fig. 2k-m**).

Overdose of the pain medicine acetaminophen (APAP) is the most common cause of drug-induced liver failure in the United States. Acute doses of acetaminophen cause hepatotoxicity^34, 35^, and the injured liver tissue enters a regenerative mode, leading to rapid healing and replacement of the damaged lobule layers^36^. However, if the doses are too high, the liver will be damaged substantially and will no longer be able to function^37, 38^. Similar to the effect on CCl4-induced injury, adenoviral LIFR delivery significantly promoted liver damage repair and regeneration in acetaminophen-treated mice (**Fig. 2n-t**). Therefore, both loss- and gain-of-function analyses demonstrate that LIFR positively regulates liver injury repair and regeneration.

### LIFR deficiency or overexpression does not affect hepatocyte proliferation *ex vivo* or *in vitro*

To determine whether Lifr regulates hepatocyte proliferation in a cell-autonomous manner, we treated primary hepatocytes isolated from *Lifr*^fl/fl^ and *Lifr*^fl/fl^;Alb-Cre mice with purified hepatocyte mitogens, HGF and EGF, which did not alter *Lifr* expression (**Extended Data Fig. 2a**). mRNA levels of cyclin D1 and PCNA (**Extended Data Fig. 2b, c**), as well as percentages of Ki67-positive cells (**Extended Data Fig. 2d, e**), were similar in the hepatocytes isolated from *Lifr*^fl/fl^ and *Lifr*^fl/fl^;Alb-Cre mice. To determine the effect of LIFR overexpression, we injected C57BL/6J mice via the tail vein with control adenovirus or LIFR-expressing adenovirus, isolated primary hepatocytes, and treated the cells with HGF and EGF. Despite the expected upregulation of proliferative markers after mitogen treatment, no significant differences were observed in cyclin D1, PCNA, and Ki67 levels between control and LIFR-overexpressing hepatocytes (**Extended Data Fig. 2f-j**). To further corroborate this result, we infected primary hepatocytes isolated from C57BL/6J mice with control or LIFR-overexpressing adenovirus *in vitro*, followed by treatment with HGF and EGF, and we found no significant changes in cyclin D1 and PCNA levels upon overexpression of LIFR (**Extended Data Fig. 2k-m**). Therefore, LIFR deficiency or overexpression does not affect hepatocyte proliferation *ex vivo* or *in vitro*, suggesting the involvement of other cell types in LIFR-regulated liver regeneration *in vivo*.

### Hepatocytic Lifr promotes neutrophil recruitment in liver regeneration models

Hepatic immune cells have important roles in liver injury repair and regeneration^7, 21, 39^. To assess the effect of hepatocyte-specific knockout of Lifr on the abundance of immune cells in injured livers, we used time-of-flight mass cytometry (CyTOF) for high-dimensional analysis of liver-infiltrating immune cells at the single-cell level^40–42^. This analysis revealed that in mice that received partial hepatectomy or CCl4 treatment, Lifr deficiency significantly decreased the abundance of hepatic neutrophils (**Fig. 3a-d**), without affecting the abundance of other immune cell types in the liver (**Extended Data Fig. 3a, b**). We then performed multiplex immunofluorescence staining^42^ of liver sections with antibodies against Ly6G (a marker of neutrophils)^43, 44^ and Ki67, further confirming that compared with *Lifr*^fl/fl^ mice, *Lifr*^fl/fl^;Alb-Cre mice showed a substantial reduction in hepatic neutrophil numbers and hepatocyte proliferation levels after hepatectomy (**Fig. 3e-g**). Notably, the abundance of Ki67-positive hepatocytes correlated with the number of hepatic neutrophils in individual mice (**Fig. 3h**).

**Figure 3.**
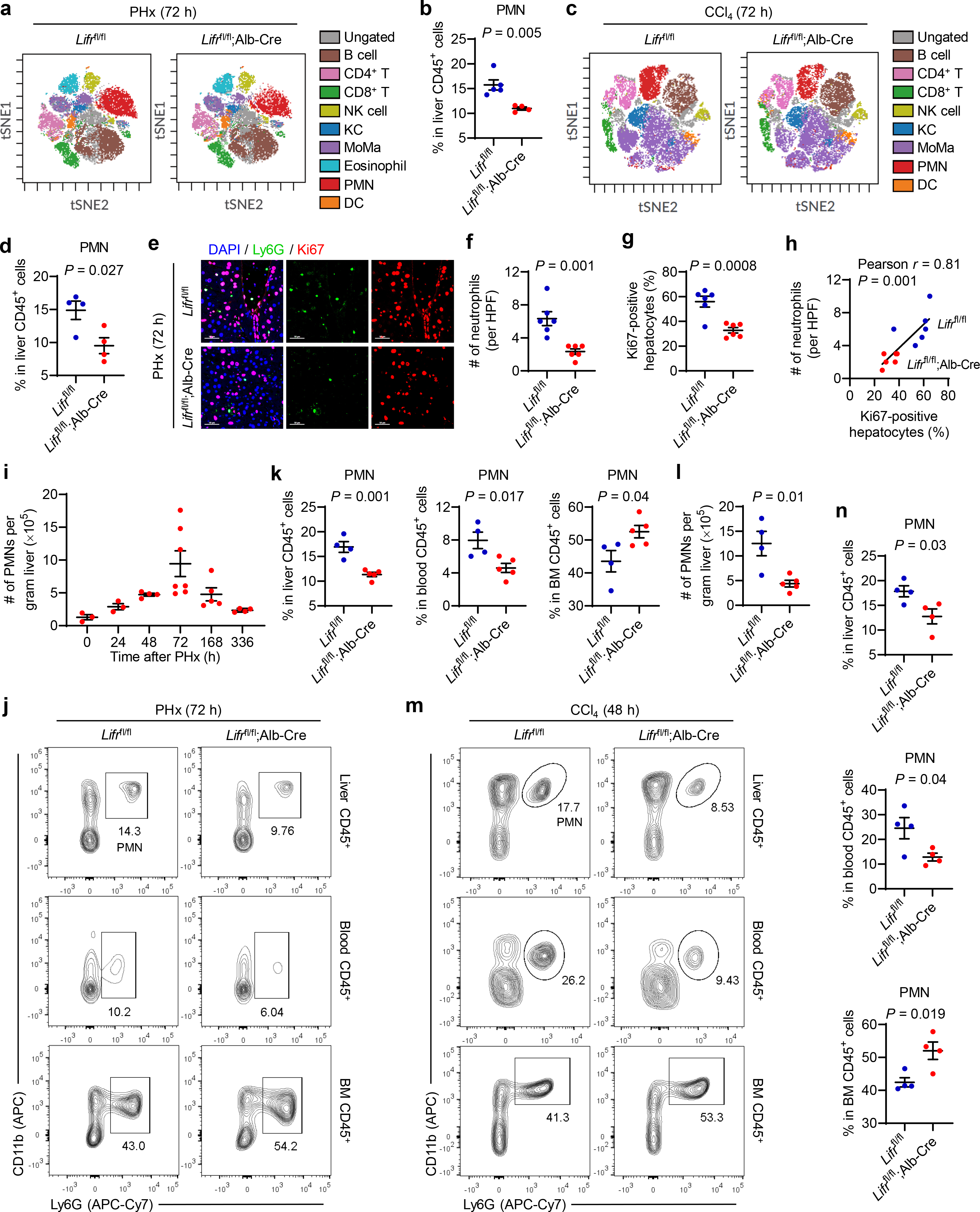
Hepatocytic Lifr promotes neutrophil recruitment in liver regeneration models. **a.** CyTOF-based immune profiling of the livers in *Lifr*^fl/fl^ and *Lifr*^fl/fl^;Alb-Cre mice at 72 hours after 2/3 partial hepatectomy (PHx). Representative viSNE plots were colored by immune cell populations. NK: natural killer cells. KC: Kupffer cells. MoMa: monocyte-derived macrophages. PMN: polymorphonuclear neutrophils. DC: dendritic cells. **b.** Quantification of liver-infiltrating PMNs in *Lifr*^fl/fl^ and *Lifr*^fl/fl^;Alb-Cre mice at 72 hours after PHx. *n* = 5 and 4 mice. **c.** CyTOF-based immune profiling of the livers in *Lifr*^fl/fl^ and *Lifr*^fl/fl^;Alb-Cre mice at 72 hours after CCl4 treatment. Representative viSNE plots were colored by immune cell populations. **d.** Quantification of liver-infiltrating PMNs in *Lifr*^fl/fl^ and *Lifr*^fl/fl^;Alb-Cre mice at 72 hours after CCl4 treatment. *n* = 4 mice per group. **e-h.** Multiplex immunofluorescence staining of Ly6G (green) and Ki67 (red) on liver sections (**e**), the number of neutrophils per high-power field (HPF; **f**), percentage of Ki67-positive hepatocytes (**g**), and correlation between hepatic neutrophil abundance and Ki67-positive hepatocyte abundance (**h**) at 72 hours after PHx. Scale bars, 50 μm. *n* = 6 mice per group. Statistical significance in **h** was determined by the Pearson correlation test. **i.** Number of PMNs per gram of liver in C57BL/6J mice at the indicated time points after PHx. *N* = 3, 3, 4, 7, 5, and 4 mice. **j, k.** Representative flow cytometry plots (**j**) and percentage of neutrophils (**k**) in liver, blood, and bone marrow (BM) CD45^+^ cells from *Lifr*^fl/fl^ and *Lifr*^fl/fl^;Alb-Cre mice at 72 hours after PHx. *n* = 4 and 5 mice. **l.** Number of PMNs per gram of liver in the mice described in **j**. *n* = 4 and 5 mice. **m, n.** Representative flow cytometry plots (**m**) and percentage of neutrophils (**n**) in liver, blood, and bone marrow (BM) CD45^+^ cells from *Lifr*^fl/fl^ and *Lifr*^fl/fl^;Alb-Cre mice at 48 hours after CCl4 treatment. *n* = 4 mice per group. Statistical significance in **b**, **d**, **f**, **g**, **k**, **l**, and **n** was determined by a two-tailed unpaired *t*-test. Error bars are s.e.m.

In response to infection or tissue damage, neutrophils are released from the bone marrow to participate in pathogen clearance or tissue repair^45–47^. Interestingly, a recent study showed that in patients who had undergone surgical resection of a diseased part of the liver, a significant increase in intrahepatic neutrophils was observed during the early response to resection^48^. Consistent with this clinical observation, we found that in mice, the number of neutrophils (CD45^+^CD11b^+^Ly6G^+^ cells, based on flow cytometry analysis) per gram of liver increased in a time-dependent manner after partial hepatectomy and peaked at 72 hours (**Fig. 3i**), validating that resection of liver tissues induces recruitment of neutrophils. Notably, compared with *Lifr*^fl/fl^ mice, *Lifr*^fl/fl^;Alb-Cre mice exhibited an increase in neutrophils in the bone marrow and a decrease in neutrophils in the blood and liver (**Fig. 3j-l**), with no changes in total hepatic CD45^+^ cells (**Extended Data Fig. 3c**). Similar results were obtained from the CCl4 model (**Fig. 3m, n**). On the other hand, adenoviral LIFR delivery led to an increase in hepatic and circulating neutrophils in mice treated with CCl4 (**Extended Data Fig. 3d-f**) or acetaminophen (**Extended Data Fig. 3g-i**). Collectively, these findings suggest that hepatocytic Lifr promotes injury-induced efflux of bone marrow neutrophils to the circulation and damaged liver, which may contribute to hepatocyte proliferation and liver regeneration.

### LIFR-mediated liver injury repair and regeneration are dependent on neutrophils

To determine whether LIFR accelerates liver regeneration through neutrophils, we depleted neutrophils with a Ly6G-specific monoclonal antibody^23, 49^ in the hepatectomy-induced liver regeneration model, in which tail vein injection of control or LIFR-expressing adenovirus was performed 10 days before partial hepatectomy (**Fig. 4a**). Six hours after the partial hepatectomy, we performed an intraperitoneal injection of isotype control (IgG) or a Ly6G-specific monoclonal antibody. Then, 48 hours after the surgery, we analyzed neutrophil abundance and the levels of liver tissue repair and functional restoration (**Fig. 4a**). By using flow cytometry-based quantification of the percentage of CD45^+^CD11b^+^Ly6G^+^ cells in total CD45^+^ cells from the liver, blood, and bone marrow, we observed that adenoviral LIFR delivery promoted injury-induced efflux of bone marrow neutrophils to the blood and liver, and we also verified the efficacy of antibody-mediated depletion of neutrophils (**Fig. 4b-g**). Relative to the control adenovirus group, mice that received LIFR-expressing adenovirus exhibited a higher liver-to-body weight ratio (**Fig. 4h**), lower serum levels of ALT and AST (**Fig. 4i, j**), and higher percentages of mitotic and Ki67- positive hepatocytes (**Fig. 4k-n**) after partial hepatectomy, and these changes were abolished by the antibody-mediated depletion of neutrophils (**Fig. 4h-n**).

**Figure 4.**
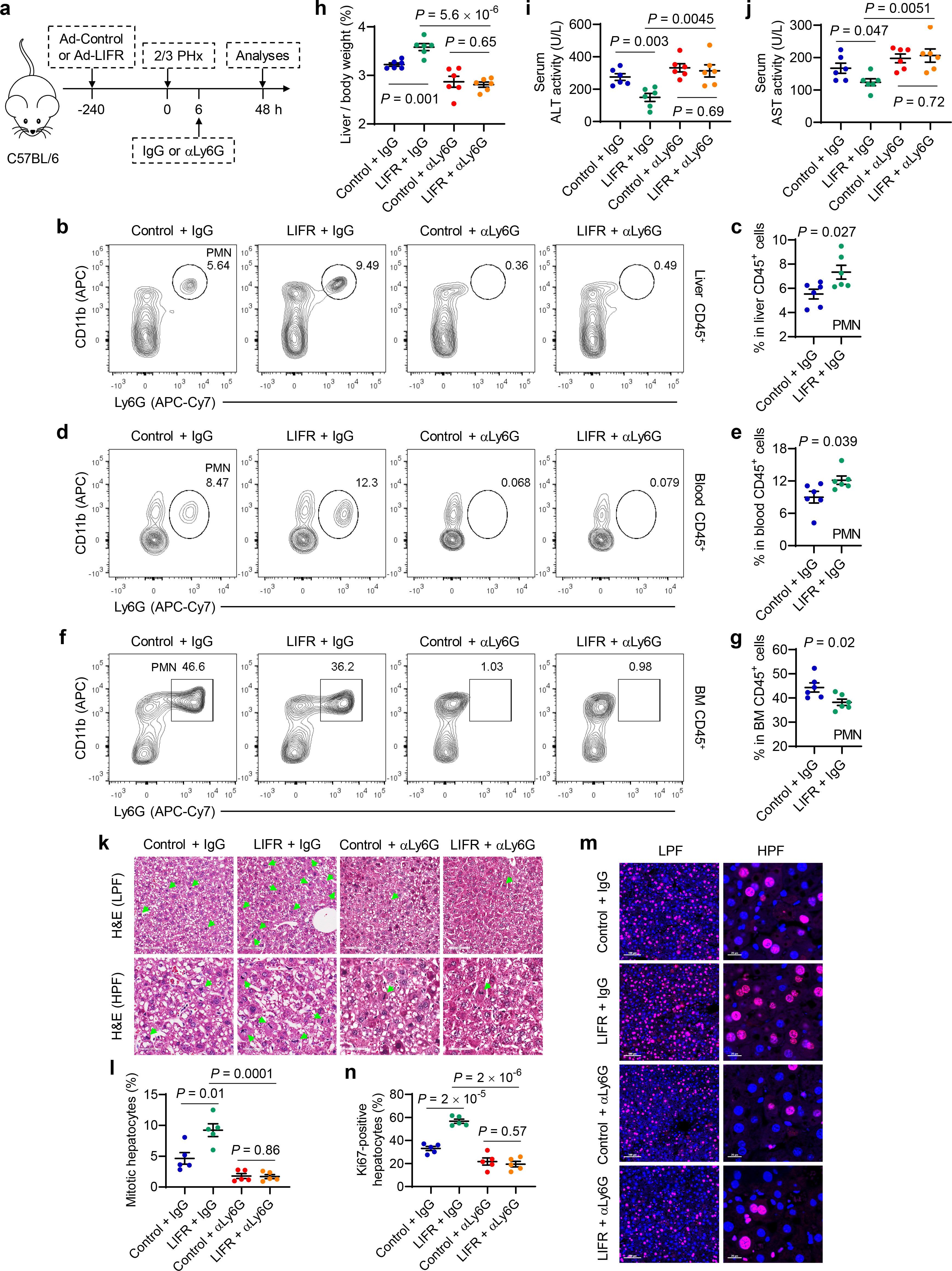
LIFR accelerates partial hepatectomy-induced liver injury repair and regeneration in a neutrophil-dependent manner. **a**-**m.** C57BL/6J mice received control or LIFR-expressing adenovirus 10 days before 2/3 partial hepatectomy (PHx). 6 hours after PHx, the mice were treated with control IgG or anti-Ly6G. Analyses were done at 48 hours after antibody treatment. **a.** Schematic of the experimental design. **b, c.** Representative flow cytometry plots (**b**) and percentage of neutrophils in liver CD45^+^ cells (**c**). *n* = 6 mice per group. **d, e.** Representative flow cytometry plots (**d**) and percentage of neutrophils in blood CD45^+^ cells (**e**). *n* = 6 mice per group. **f, g.** Representative flow cytometry plots (**f**) and percentage of neutrophils in bone marrow (BM) CD45^+^ cells (**g**). *n* = 6 mice per group. **h-j.** Liver-to-body weight ratio (**h**) and levels of serum ALT (**i**) and AST (**j**) in control and LIFR- expressing adenovirus-infected C57BL/6J mice injected with control IgG or anti-Ly6G after PHx. *n* = 6 mice per group. **k**, **l.** Representative H&E staining (**k**) and percentage of mitotic hepatocytes (**l**). Green arrows indicate mitotic nuclei. LPF: low-power field; HPF: high-power field. Scale bars, 100 μm in upper panels and 50 μm in lower panels. *n* = 5 mice per group. **m**, **n.** Immunofluorescence staining of Ki67 (**m**; overlay with DAPI staining) and percentage of Ki67-positive hepatocytes (**n**). LPF: low-power field; HPF: high-power field. Scale bars, 100 μm in left panels and 20 μm in right panels. *n* = 5 mice per group. Statistical significance in **c**, **e**, **g**, **h-j**, **l**, and **n** was determined by a two-tailed unpaired *t*-test. Error bars are s.e.m.

Similarly, in the CCl4-induced liver injury model (**Extended Data Fig. 4a**), we depleted neutrophils efficiently by using a Ly6G-specific monoclonal antibody, and we observed that LIFR overexpression led to a decrease in neutrophils in the bone marrow and an increase in neutrophils in the liver and blood (**Extended Data Fig. 4b-g**). Moreover, mice injected with LIFR-expressing adenovirus showed improved liver injury repair and organ function after CCl4 treatment, as evidenced by lower serum levels of ALT and AST (**Extended Data Fig. 4h, i**), less necrosis and TUNEL staining (**Extended Data Fig. 4j-m**), and higher percentages of mitotic and Ki67-positive hepatocytes (**Extended Data Fig. 4n, o**) after partial hepatectomy, and these changes were abrogated by neutrophil depletion (**Extended Data Fig. 4h-o**). We conclude from these experiments that hepatocytic LIFR accelerates surgery- and chemical-induced liver injury repair and regeneration in a neutrophil-dependent manner.

### Hepatocytic Lifr promotes HGF production through neutrophils

Hepatocyte proliferation during liver regeneration involves extracellular factors; two of these factors, HGF and EGF, are complete mitogens, and other factors, e.g., IL-6, TNF-α, TGF-β, are auxiliary mitogens^2, 18, 50, 51^. To investigate whether these hepatocyte mitogens are involved in the role of LIFR in regulating liver regeneration, we measured the levels of HGF, EGF, IL-6, TNF-α, and TGF-β in the serum of *Lifr*^fl/fl^ and *Lifr*^fl/fl^;Alb-Cre mice after partial hepatectomy or CCl4 treatment. We found that HGF levels were considerably lower in *Lifr*^fl/fl^;Alb-Cre mice compared with *Lifr*^fl/fl^ mice, whereas no significant difference was observed in serum levels of EGF, IL-6, TNF-α, and TGF-β between these two groups (**Fig. 5a-c**). A recent clinical investigation revealed that patients who underwent partial hepatectomy had a significant increase in intrahepatic neutrophils^48^. In our study, injection of C57BL/6J mice with LIFR-expressing adenovirus led to elevated levels of serum HGF after partial hepatectomy, which was reversed by neutrophil depletion with a Ly6G-specific monoclonal antibody^23^ (**Fig. 5d**), suggesting that LIFR increases HGF production in a neutrophil-dependent manner.

**Figure 5.**
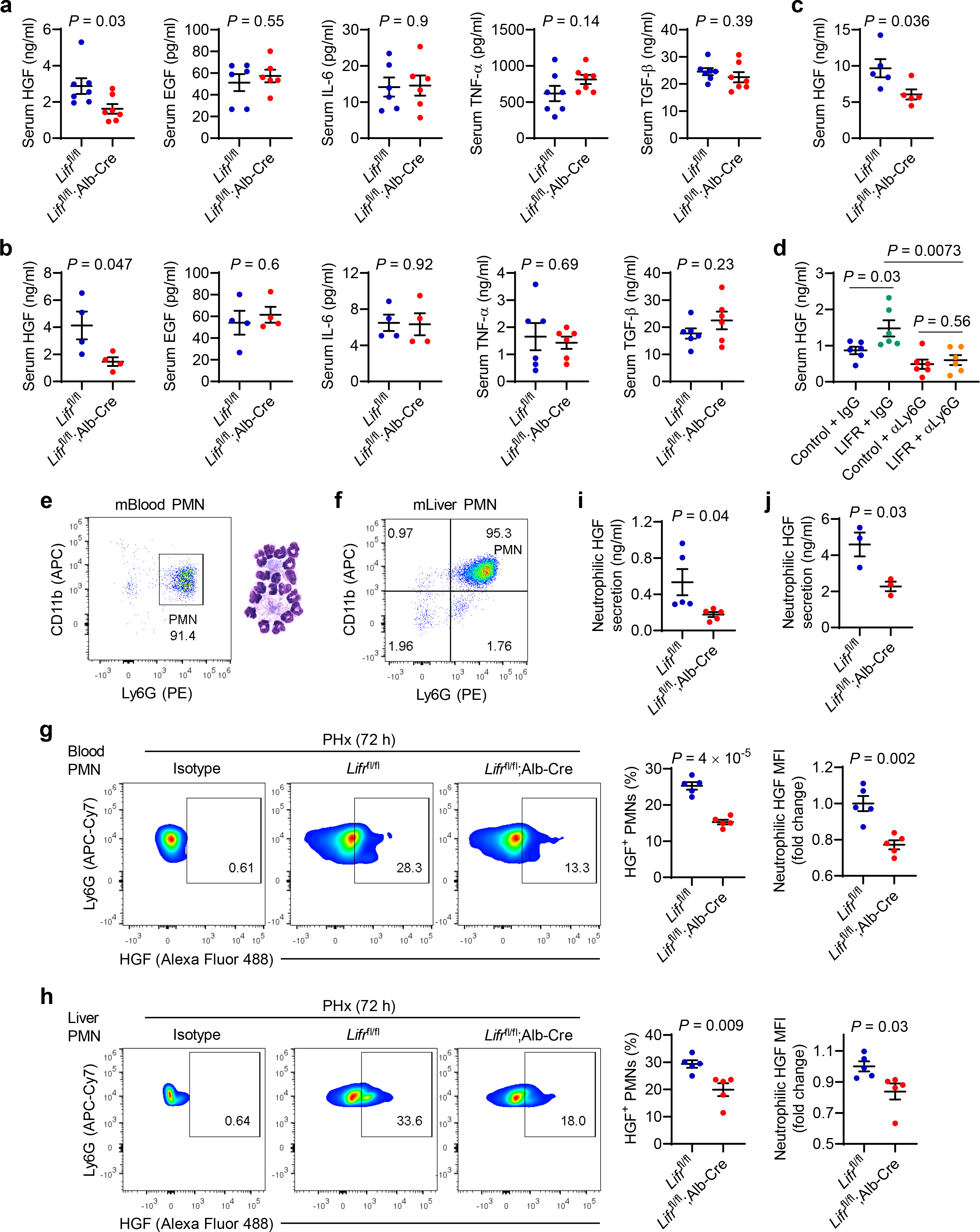
Hepatocytic Lifr promotes HGF production through neutrophils. **a.** Serum HGF, EGF, IL-6, TNF-α, and TGF-β levels in *Lifr*^fl/fl^ and *Lifr*^fl/fl^;Alb-Cre mice at 72 hours after 2/3 partial hepatectomy (PHx). *n* = 7, 7, 6, 6, 6, 6, 7, 7, 7, and 7 mice. **b.** Serum HGF, EGF, IL-6, TNF-α, and TGF-β levels in *Lifr*^fl/fl^ and *Lifr*^fl/fl^;Alb-Cre mice at 72 hours after CCl4 treatment. *n* = 4, 4, 4, 4, 4, 4, 6, 6, 6, and 6 mice. **c.** Serum HGF levels in *Lifr*^fl/fl^ and *Lifr*^fl/fl^;Alb-Cre mice at 48 hours after CCl4 treatment. *n* = 5 mice per group. **d.** Serum HGF levels in control and LIFR-expressing adenovirus-infected C57BL/6J mice injected with control IgG or anti-Ly6G 48 hours after PHx. Adenovirus was administered 10 days before PHx. *n* = 6 mice per group. **e.** Flow cytometry analysis and Giemsa staining of neutrophils purified from mouse blood. **f.** Flow cytometry analysis of neutrophils purified from mouse livers. **g.** Representative flow cytometry plots, percentage of HGF^+^ PMNs, and neutrophilic HGF staining (quantified by mean fluorescence intensity, MFI) in blood neutrophils from *Lifr*^fl/fl^ and *Lifr*^fl/fl^;Alb- Cre mice at 72 hours after PHx. *n* = 5 mice per group. **h.** Representative flow cytometry plots, percentage of HGF^+^ PMNs, and neutrophilic HGF staining in liver-infiltrating neutrophils from *Lifr*^fl/fl^ and *Lifr*^fl/fl^;Alb-Cre mice at 72 hours after PHx. *n* = 5 mice per group. **i.** ELISA of HGF in the conditioned medium of purified blood neutrophils. *n* = 5 mice per group. **j.** ELISA of HGF in the conditioned medium of purified liver-infiltrating neutrophils. *n* = 3 mice per group. Statistical significance in **a-d** and **g-j** was determined by a two-tailed unpaired *t*-test. Error bars are s.e.m.

To further corroborate that neutrophils are a Lifr-regulated source of HGF, we isolated polymorphonuclear neutrophils from mouse blood and livers (**Fig. 5e, f**). Notably, at 72 hours after partial hepatectomy, *Lifr*^fl/fl^;Alb-Cre mice had much lower percentages of HGF-positive neutrophils in the blood and liver than did *Lifr*^fl/fl^ mice (**Fig. 5g, h**). Moreover, knockout of Lifr in hepatocytes reduced the *ex vivo* production of HGF by neutrophils isolated from either the blood or the liver at 72 hours after partial hepatectomy (**Fig. 5i, j**). Taken together, our results indicate that hepatocytic Lifr promotes the production of the pro-regenerative growth factor HGF through neutrophils.

### LIFR upregulates hepatocyte-derived cholesterol, which acts on neutrophils to boost HGF production

The liver is the central organ that controls lipid homeostasis^52^. Considering that lipids are regulators of neutrophil viability, recruitment, and functions^53^, we performed a lipidomic analysis of plasma samples collected from *Lifr*^fl/fl^ and *Lifr*^fl/fl^;Alb-Cre mice at 72 hours after partial hepatectomy (**Fig. 6a**). The global lipid profile showed that phosphatidylinositol, acylcarnitine, cholesterol, cholesterol ester, and phosphatidylcholine were downregulated, whereas diacylglycerol and triacylglycerol were upregulated in *Lifr*^fl/fl^;Alb-Cre mice (**Fig. 6b-d** and **Supplementary Table 1**).

**Figure 6.**
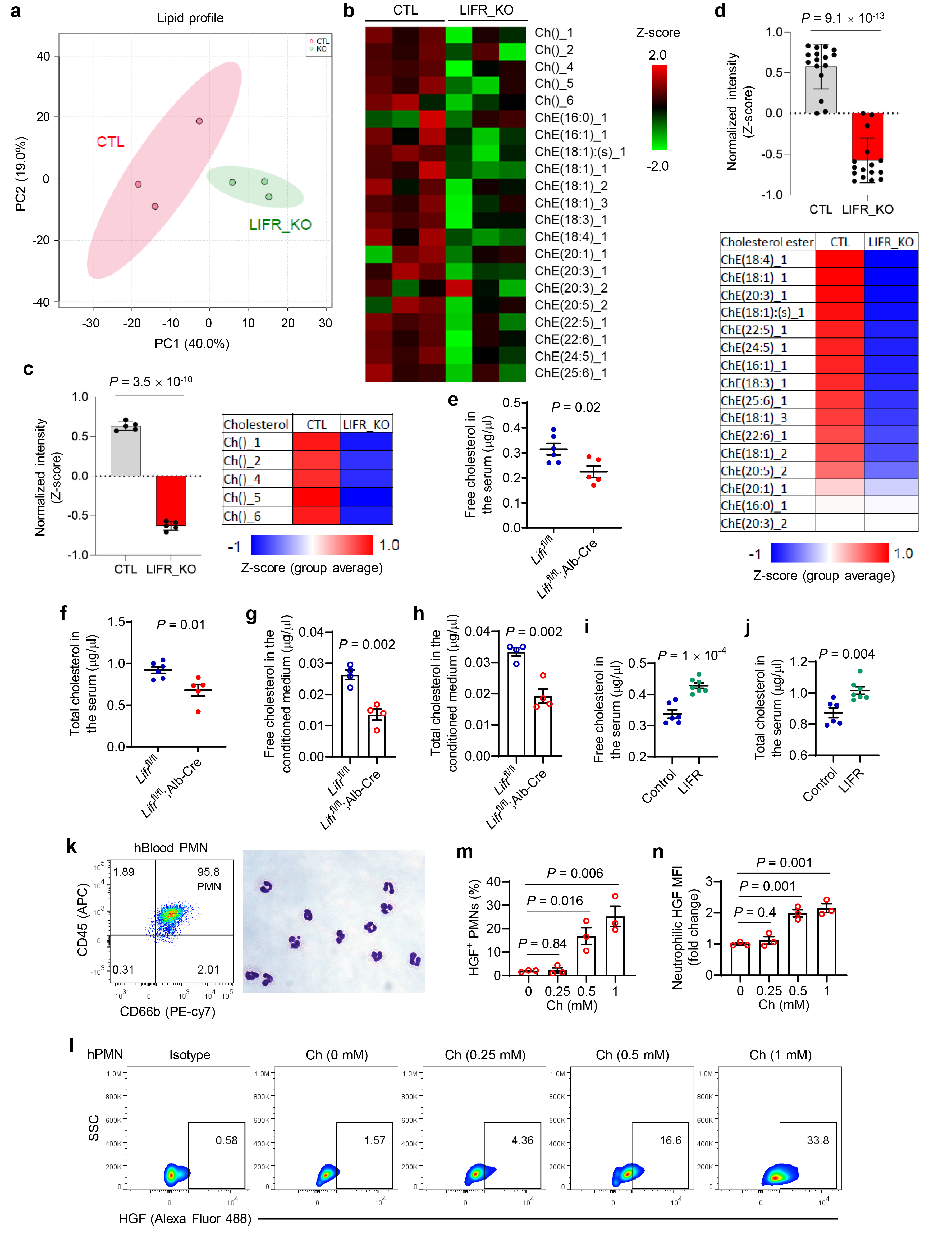
LIFR upregulates hepatocyte-derived cholesterol, which acts on neutrophils to boost HGF production. **a-d.** Lipidomic analysis of plasma samples collected from *Lifr*^fl/fl^ (CTL) and *Lifr*^fl/fl^;Alb-Cre (LIFR_KO) mice at 72 hours after 2/3 partial hepatectomy (PHx). *n* = 3 mice per group. **a.** Lipid profile presented as a Principal Component Analysis plot. **b.** Heatmap of plasma cholesterol (Ch) and cholesterol ester (ChE) levels. **c.** Plasma cholesterol (Ch) levels in *Lifr*^fl/fl^ and *Lifr*^fl/fl^;Alb-Cre mice. **d.** Plasma cholesterol ester (ChE) levels in *Lifr*^fl/fl^ and *Lifr*^fl/fl^;Alb-Cre mice. **e.** Free cholesterol levels in the serum of *Lifr*^fl/fl^ and *Lifr*^fl/fl^;Alb-Cre mice at 72 hours after PHx. *N* = 6 and 5 mice. **f.** Total cholesterol levels in the serum of *Lifr*^fl/fl^ and *Lifr*^fl/fl^;Alb-Cre mice at 72 hours after PHx. *N* = 6 and 5 mice. **g.** Free cholesterol levels in the conditioned medium of primary hepatocytes isolated from *Lifr*^fl/fl^ and *Lifr*^fl/fl^;Alb-Cre mice. *n* = 4 biological replicates per group. **h.** Total cholesterol levels in the conditioned medium of primary hepatocytes isolated from *Lifr*^fl/fl^ and *Lifr*^fl/fl^;Alb-Cre mice. *n* = 4 biological replicates per group. **i.** Free cholesterol levels in the serum of control and LIFR-expressing adenovirus-injected C57BL/6J mice at 48 hours after PHx. *n* = 6 and 7 mice. **j.** Total cholesterol levels in the serum of control and LIFR-expressing adenovirus-injected C57BL/6J mice at 48 hours after PHx. *n* = 6 and 7 mice. **k.** Flow cytometry and Giemsa staining of neutrophils purified from human blood. **l-n.** Representative flow cytometry plots (**l**), percentage of HGF^+^ PMNs (**m**), and quantification of neutrophilic HGF (by mean fluorescence intensity, MFI; **n**) in human neutrophils treated with 0, 0.25, 0.5, or 1 mM cholesterol (Ch) for 4 hours. *n* = 3 biological replicates per group. Statistical significance in **c-j**, **m**, and **n** was determined by a two-tailed unpaired *t*-test. Error bars are s.e.m.

Of note, cholesterol has previously been shown to modulate neutrophil functions^53, 54^ and liver regeneration^55^. To validate our lipidomics results, we measured levels of free and total cholesterol in the serum samples collected from *Lifr*^fl/fl^ and *Lifr*^fl/fl^;Alb-Cre mice by using a cholesterol quantitation kit, finding that levels of both free and total cholesterol were reduced in *Lifr*^fl/fl^;Alb- Cre mice compared with *Lifr*^fl/fl^ mice after partial hepatectomy (**Fig. 6e, f**). To verify that this difference was attributable to hepatocyte-derived cholesterol, we isolated primary hepatocytes from *Lifr*^fl/fl^ and *Lifr*^fl/fl^;Alb-Cre mice, and measured cholesterol levels in the conditioned medium. We found that Lifr deficiency reduced the *ex vivo* production of cholesterol by hepatocytes (**Fig. 6g, h**). On the other hand, adenoviral LIFR delivery resulted in higher levels of serum cholesterol after liver injury (**Fig. 6i, j**). To determine whether cholesterol regulates levels of neutrophil- derived HGF, we isolated polymorphonuclear neutrophils from human blood (**Fig. 6k**) and treated them with 0.25-1 mM cholesterol for 4 hours, finding that cholesterol treatment elevated HGF production in a dose-dependent manner (**Fig. 6l-n**). We conclude that LIFR upregulates hepatocyte-derived cholesterol, which acts on neutrophils to boost HGF production.

### Hepatocytic Lifr promotes neutrophil recruitment through Cxcl1

Chemokines are a family of small secreted proteins that act as attractants for different types of leukocytes, including neutrophils, and recruit them to sites of infection, inflammation, and injury^56^. Neutrophil recruitment is mainly mediated by chemokines that have a glutamate-leucine-arginine motif (ELR chemokines). ELR members, C-X-C motif ligand 1 (CXCL1), CXCL2, and CXCL5, bind to C-X-C motif chemokine receptor 2 (CXCR2), a major chemokine receptor expressed by neutrophils in both mice and humans, which is responsible for the mobilization of neutrophils from the bone marrow^47, 57–59^. In addition, CXCL12-CXCR4 signaling plays a pivotal role in the retention of neutrophils in the bone marrow and homing of senescent neutrophils back to the bone marrow^60–62^. The opposing effects of CXCR2 and CXCR4 on neutrophils maintain the homeostasis of neutrophils in circulation^47, 61^. To determine whether chemokine signaling is involved in the role of hepatocytic Lifr in neutrophil recruitment in liver regeneration models, we measured *Cxcl1*, *Cxcl2*, and *Cxcl5* mRNA levels in the livers of *Lifr*^fl/fl^ and *Lifr*^fl/fl^;Alb-Cre mice after 2/3 partial hepatectomy or CCl4 treatment. Notably, hepatic *Cxcl1* levels were markedly upregulated after surgery, and this upregulation was diminished by knockout of Lifr (**Fig. 7a**). These changes were further validated by ELISA of serum levels of Cxcl1 protein in *Lifr*^fl/fl^ and *Lifr*^fl/fl^;Alb-Cre mice (**Fig. 7b**). In contrast, levels of *Cxcl2* and *Cxcl5* showed no significant changes after resection and were not affected by Lifr deficiency (**Fig. 7c, d**). Similar results were also obtained in the CCl4- induced injury model (**Fig. 7e-g**).

**Figure 7.**
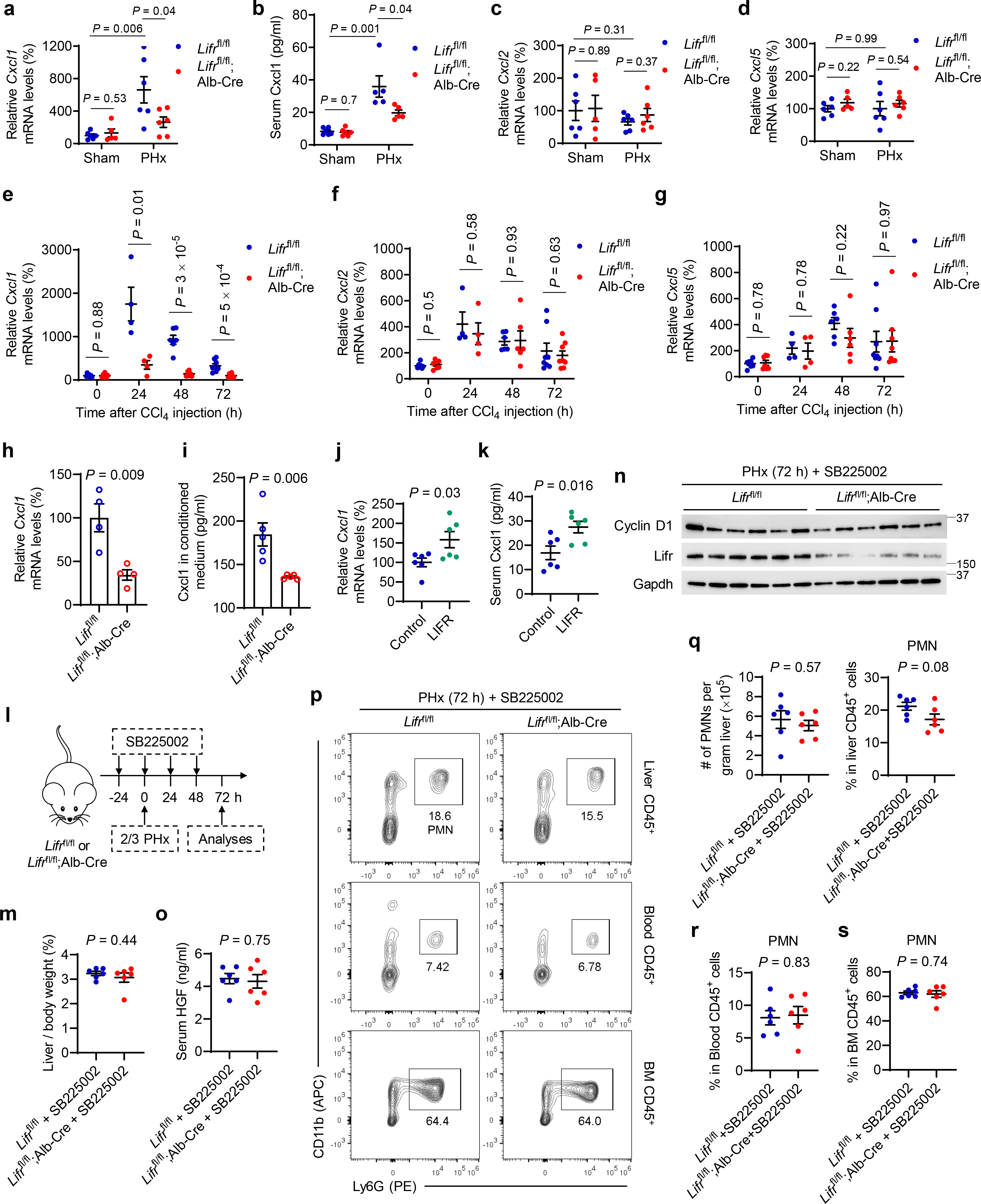
Hepatocytic Lifr promotes neutrophil recruitment through Cxcl1. **a.** qPCR of *Cxcl1* in the livers of *Lifr*^fl/fl^ and *Lifr*^fl/fl^;Alb-Cre mice at 72 hours after sham surgery or 2/3 partial hepatectomy (PHx). *n* = 6, 5, 6, and 6 mice. **b.** Serum Cxcl1 levels in *Lifr*^fl/fl^ and *Lifr*^fl/fl^;Alb-Cre mice at 72 hours after sham surgery or PHx. *n* = 6, 6, 5, and 5 mice. **c, d.** qPCR of *Cxcl2* (**c**), and *Cxcl5* (**d**) in the livers of *Lifr*^fl/fl^ and *Lifr*^fl/fl^;Alb-Cre mice at 72 hours after sham surgery or PHx. *n* = 6, 5, 6, and 6 mice. **e-g.** qPCR of *Cxcl1* (**e**)*, Cxcl2* (**f**), and *Cxcl5* (**g**) in the livers of *Lifr*^fl/fl^ and *Lifr*^fl/fl^;Alb-Cre mice at the indicated times after CCl4 treatment. *n* = 7, 7, 4, 4, 6, 6, 8, and 8 mice. **h.** qPCR of *Cxcl1* in primary hepatocytes isolated from *Lifr*^fl/fl^ and *Lifr*^fl/fl^;Alb-Cre mice. *n* = 4 biological replicates per group. **i.** ELISA of Cxcl1 in the conditioned medium of primary hepatocytes isolated from *Lifr*^fl/fl^ and *Lifr*^fl/fl^;Alb-Cre mice. *n* = 5 biological replicates per group. **j.** qPCR of *Cxcl1* in the livers of control and LIFR-expressing adenovirus-injected C57BL/6J mice at 48 hours after PHx. *n* = 6 mice per group. **k.** Serum Cxcl1 levels in control and LIFR-expressing adenovirus-injected C57BL/6J mice at 48 hours after PHx. *n* = 6 mice per group. **l-s.** *Lifr*^fl/fl^ and *Lifr*^fl/fl^; Alb-Cre mice underwent PHx and received SB225002 treatment for 4 days. **l.** Schematic of the experimental design. **m.** Liver-to-body weight ratio at 72 hours after PHx. *n* = 6 mice per group. **n.** Immunoblotting of Lifr, cyclin D1, and Gapdh in the livers. **o.** Serum HGF levels. *n* = 6 mice per group. **p-s.** Representative flow cytometry plots (**p**) and quantification of neutrophils in liver (**q**), blood (**r**), and bone marrow (BM; **s**) CD45^+^ cells from *Lifr*^fl/fl^ and *Lifr*^fl/fl^;Alb-Cre mice at 72 hours after PHx. *n* = 6 mice per group. Statistical significance in **a-k**, **m**, **o**, and **q-s** was determined by a two-tailed unpaired *t*-test. Error bars are s.e.m.

To determine whether knockout of Lifr downregulates hepatocyte-secreted Cxcl1, we isolated primary hepatocytes from *Lifr*^fl/fl^ and *Lifr*^fl/fl^;Alb-Cre mice and collected the conditioned medium. We found that hepatocytic Lifr deficiency reduced both *Cxcl1* mRNA levels as well as the *ex vivo* production of Cxcl1 by hepatocytes (**Fig. 7h, i**). On the other hand, adenoviral LIFR delivery led to upregulation of both hepatic *Cxcl1* mRNA and serum Cxcl1 protein after liver injury (**Fig. 7j, k**). Therefore, upon liver damage, LIFR promotes Cxcl1 production in hepatocytes.

To determine whether blockade of Cxcl1-Cxcr2 signaling abrogates the observed effects of Lifr loss on liver regeneration, we treated *Lifr*^fl/fl^ and *Lifr*^fl/fl^;Alb-Cre mice with a CXCR2 antagonist, SB225002^63–65^ (**Fig. 7l**). Unlike untreated mice (**Extended Data Fig. 1c**; **Fig. 1b**, **3j-l**, and **5a**), *Lifr*^fl/fl^ and *Lifr*^fl/fl^;Alb-Cre mice treated with SB225002 showed no significant difference in the liver-to-body weight ratio (**Fig. 7m**), cyclin D1 expression (**Fig. 7n**), serum HGF levels (**Fig. 7o**), or the abundance of polymorphonuclear neutrophils in the liver, blood, and bone marrow (**Fig. 7p- s**) after partial hepatectomy, suggesting that Cxcl1-Cxcr2 signaling is required for the observed loss-of-function effects of Lifr.

### LIFR regulates CXCL1 through STAT3, and treatment with a STAT3 inhibitor reverses LIFR-accelerated liver regeneration

LIFR can regulate gp130-JAK-STAT3, MAPK, PI3K, Hippo-YAP, and NF-κB pathways in tissue-dependent and context-dependent manners^29, 66–71^. Recently, we reported that in liver cancer models, knockout of Lifr promoted tumorigenesis and conferred resistance to sorafenib-induced ferroptosis through NF-κB-dependent upregulation of the iron-sequestering protein lipocalin-2 (Lcn2)^29^. However, although lipocalin-2 is upregulated after partial hepatectomy, Lcn2-deficient mice and wild-type mice showed no difference in surgical injury-induced hepatocyte proliferation and liver regeneration^72^, suggesting that Lcn2 is not a regulator of the hepatic regeneration process. On the other hand, the gp130-Stat3 pathway has been implicated in liver regeneration in mice^73–76^. Moreover, Cxcl1 expression is induced by the gp130-Stat3 axis in mouse models of infection and inflammation^65, 77^. Therefore, we hypothesized that LIFR activates gp130-STAT3 signaling in hepatocytes after liver injury, which in turn induces CXCL1 expression to recruit reparative neutrophils. Indeed, 48 hours after partial hepatectomy, *Lifr*^fl/fl^;Alb-Cre mice showed much less Stat3 phosphorylation in their livers compared with *Lifr*^fl/fl^ mice (**Fig. 8a**); on the other hand, adenoviral delivery of LIFR elevated phospho-Stat3 levels in the mouse liver (**Fig. 8b**).

**Figure 8.**
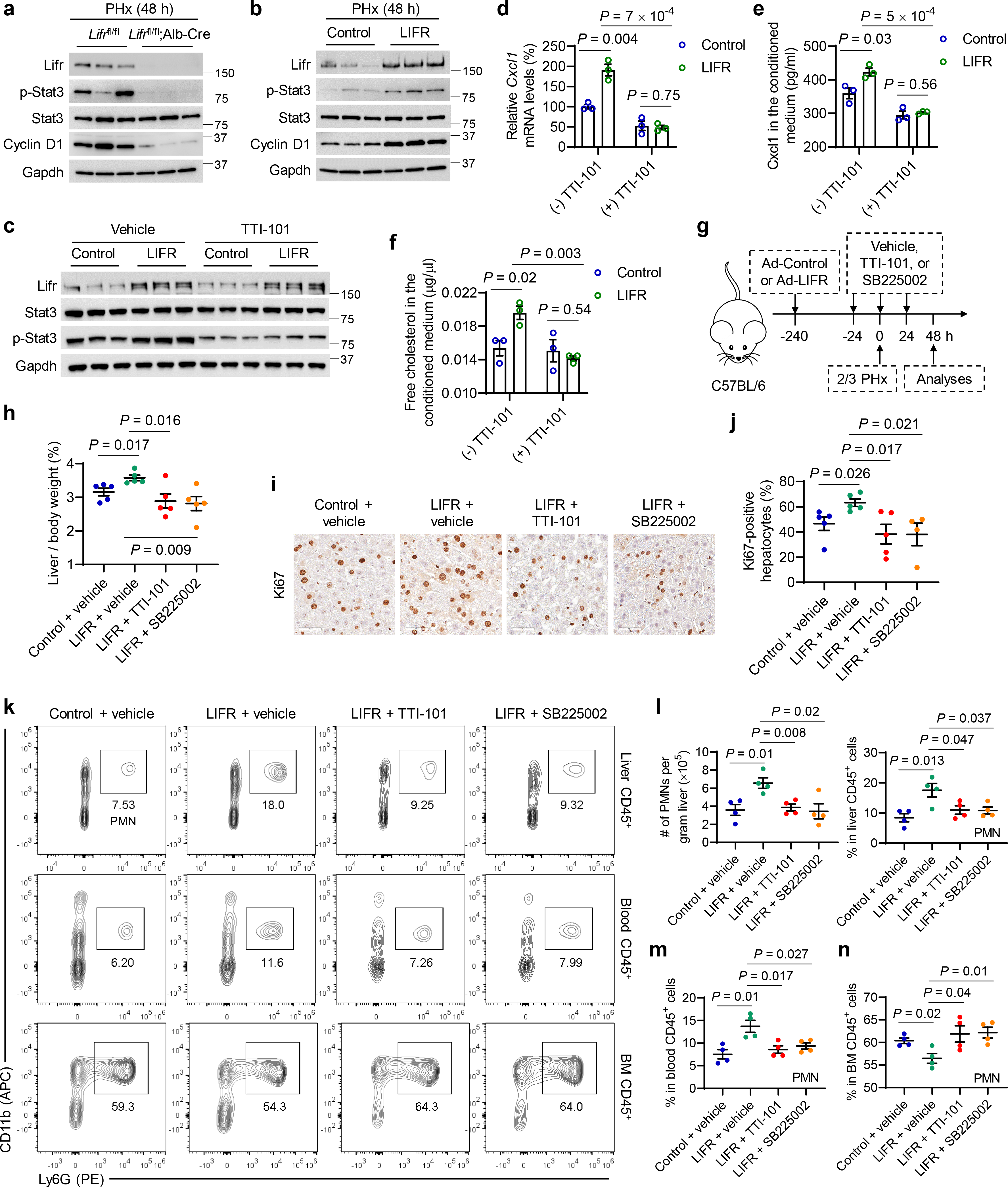
LIFR regulates CXCL1 through STAT3, and treatment with a STAT3 inhibitor reverses LIFR-accelerated liver regeneration. **a.** Immunoblotting of Lifr, Stat3, p-Stat3, cyclin D1, and Gapdh in the livers of *Lifr*^fl/fl^ and *Lifr*^fl/fl^;Alb-Cre mice 48 hours after 2/3 partial hepatectomy (PHx). **b.** Immunoblotting of Lifr, Stat3, p-Stat3, cyclin D1, and Gapdh in the livers of control and LIFR- expressing adenovirus-injected C57BL/6J mice 48 hours after PHx. **c.** Immunoblotting of Lifr, Stat3, p-Stat3, and Gapdh in primary hepatocytes isolated from C57BL/6J mice. Hepatocytes were treated with10 µM TTI-101 or vehicle for 72 hours and infected with control or LIFR-expressing adenovirus for 24 hours. **d.** qPCR of *Cxcl1* in the hepatocytes described in **c**. *n* = 3 biological replicates per group. **e, f.** Levels of secreted Cxcl1 (**e**) and free cholesterol (**f**) in the conditioned medium of the hepatocytes described in **c**. *n* = 3 biological replicates per group. **g**-**n.** C57BL/6J mice received control or LIFR-expressing adenovirus 10 days before PHx and were treated with TTI-101 or SB225002 for 3 days. Analyses were done at 48 hours after PHx. **g.** Schematic of the experimental design. **h.** Liver-to-body weight ratio at 48 hours after PHx. *n* = 5 mice per group. **i**, **j.** Immunohistochemical staining of Ki67 of liver sections (**i**) and percentage of Ki67-positive hepatocytes (**j**) at 48 hours after PHx. Scale bars, 50 μm. *n* = 5, 5, 5, and 4 mice. **k**-**n.** Representative flow cytometry plots (**k**) and quantification of neutrophils in liver (**l**), blood (**m**), and bone marrow (BM) (**n**) CD45^+^ cells at 48 hours after PHx. *n* = 4 mice per group. Statistical significance in **d-f**, **h**, **j**, and **l-n** was determined by a two-tailed unpaired *t*-test. Error bars are s.e.m.

Next, we isolated primary hepatocytes from C57BL/6J mice, treated these cells with a STAT3 inhibitor TTI-101 (also known as C188-9)^78–80^, and infected them with control or LIFR-expressing adenovirus. As expected, overexpression of LIFR in hepatocytes upregulated Stat3 phosphorylation, Cxcl1 production, and cholesterol secretion, which was abolished by TTI-101 treatment (**Fig. 8c-f**). Finally, to determine whether LIFR accelerates liver regeneration through STAT3-CXCL1-CXCR2 signaling, we injected control or LIFR-expressing adenovirus into C57BL/6J mice 10 days before partial hepatectomy and then treated these mice with TTI-101 or SB225002 for 3 days (**Fig. 8g**). As expected, LIFR overexpression promoted liver regeneration and hepatocyte proliferation after injury, which was abolished by treatment with TTI-101 or SB225002 (**Fig. 8h-j**). TTI-101 treatment also dampened LIFR-induced production of Cxcl1, cholesterol, and HGF (**Extended Data Fig. 5a-e**). Moreover, STAT3 inhibition or CXCR2 blockade abrogated LIFR-induced efflux of bone marrow neutrophils to the blood and injured liver after hepatectomy (**Fig. 8k-n**). Altogether, these findings suggest that STAT3-CXCL1-CXCR2 signaling mediates, at least in part, LIFR-induced neutrophil recruitment, HGF production, and liver regeneration.

## Discussion

The link between organ regeneration and cancer is complex. On the one hand, excessive regeneration may potentially induce tumorigenesis due to overgrowth. On the other hand, deficiency in regeneration may also promote cancer due to tissue damage that cannot be repaired; however, the molecular link between impairment of tissue repair/regeneration and induction of tumorigenesis remains elusive. Intriguingly, the expression of LIFR is frequently downregulated in liver cancer but is upregulated during the recovery phase after liver injury. Previously, we found that loss of Lifr in mouse hepatocytes promoted liver tumorigenesis^29^. In the present study, hepatocyte-specific ablation of Lifr in mice delayed, whereas adenovirus-mediated delivery of LIFR accelerated liver injury repair and regeneration. Taken together, our findings reveal the role of a liver tumor suppressor in liver regeneration and a potential molecular link between impaired tissue repair and cancer.

Interestingly, LIFR deficiency or overexpression does not affect hepatocyte proliferation *ex vivo* or *in vitro*, suggesting the involvement of other cell types in LIFR-accelerated liver regeneration *in vivo*. Of note, our CyTOF-based immune profiling analysis showed that in mice that received partial hepatectomy or CCl4 treatment, hepatocytic Lifr deficiency reduced the abundance of liver- infiltrating neutrophils without affecting the abundance of other hepatic immune cell types. Notably, the involvement of neutrophils in tissue injury repair has been increasingly recognized^23–27^. After partial hepatectomy or chemical damage (by CCl4 or acetaminophen treatment) to the liver, neutrophils accumulate at the injury site and actively participate in clearing dead cells and stimulating hepatocyte proliferation, liver tissue repair, and regeneration^23, 24, 48^. Importantly, a recent study reported that genetic ablation of neutrophils in mice (by knockout of *Gcsf*) exacerbated tissue damage and diminished liver regeneration after injury, demonstrating the functional contribution of neutrophils to liver injury repair^23^. However, it has been unclear how the damaged liver signals to the bone marrow to release neutrophils to the bloodstream and liver, and how these neutrophils signal to hepatocytes to re-enter the cell cycle and proliferate. Here, we show that in response to liver damage, hepatocytic LIFR induces the secretion of the neutrophil chemoattractant CXCL1 and cholesterol in a STAT3-dependent manner. Cholesterol, in turn, acts on the recruited neutrophils to secrete HGF to accelerate liver injury repair and regeneration (**Extended Data Fig. 5f**). This may have clinical implications; for instance, it would be interesting to explore whether a short course of a cholesterol-supplemented diet could help the liver to regenerate in patients who have undergone surgical resection of a diseased part of the liver.

Future studies should address the following issues. First, we found that Lifr expression is upregulated during the recovery phase after liver damage; yet, how this receptor is regulated under hepatic injury conditions is unknown. Second, our study identified a LIFR-STAT3-CXCL1- CXCR2 axis that promotes injury-induced efflux of neutrophils from the bone marrow to the circulation and damaged liver, but additional factors could also be involved in neutrophil recruitment. Third, this study identified a LIFR-STAT3-cholesterol-HGF axis that mediates hepatic damage-induced crosstalk between hepatocytes and neutrophils to stimulate hepatocyte proliferation, but how cholesterol acts on neutrophils to boost HGF production remains an open question. Finally, our previous study^29^ and the present study collectively revealed liver-intrinsic mechanisms of LIFR in suppressing HCC and liver-extrinsic mechanisms of LIFR in promoting liver injury repair and regeneration. Whether LIFR has similar or different functions and mechanisms of action in other organs warrants further investigation with *in vivo* models.

## Methods

### Mice

All animal studies were performed in accordance with a protocol approved by the Institutional Animal Care and Use Committee of MD Anderson Cancer Center. Animals were housed at 70 °F- 74 °F (set point: 72 °F) with 40%-55% humidity (set point: 45%). The light cycle in the animal rooms is 12 hours of light and 12 hours of dark. C57BL/6J mice were from MD Anderson’s internal supply or the Jackson Laboratory (JAX: 000664). The generation of *Lifr*^flox/flox^ (*Lifr*^fl/fl^) mice and hepatocyte-specific *Lifr* deletion mutants (*Lifr*^fl/fl^;Alb-Cre) and primers for PCR genotyping were described in our previous publication^29^.

### Partial hepatectomy and drug treatment

The 2/3 partial hepatectomy (PHx) surgery was described previously^31^. We performed the surgery when the mice were 8-12 weeks old. Briefly, after the upper abdomen was shaved and the skin was disinfected, mice were anesthetized with isoflurane. A medium incision was made on the upper abdomen to expose the liver. The left lateral lobe and the median lobe were surgically removed while mice were under isoflurane anesthesia. After the resection of lobes, the peritoneum was closed with absorbable sutures (Oasis, MV-J397-V), and the skin was closed using the BD AutoClip Wound Closing System (BD, 427630). All animal experiments took place in a specific pathogen-free facility of MD Anderson Cancer Center. For drug treatment, the CXCR2 antagonist SB225002^63–65^ (Selleckchem, S7651) was dissolved in 5% dimethylsulfoxide (DMSO) in olive oil (Sigma-Aldrich, O1514) and injected intraperitoneally at a concentration of 2 mg/kg body weight at the indicated times. The STAT3 inhibitor TTI-101^81, 82^ (Selleckchem, S8605) was dissolved in 5% DMSO in olive oil (Sigma-Aldrich, O1514) and injected intraperitoneally at a concentration of 50 mg/kg body weight at the indicated times. For neutrophil depletion, mice were injected intraperitoneally with 200 μg anti-Ly6G^23, 43, 49^ (Bio X Cell, BE0075-1, clone 1A8; RRID: AB_1107721) or rat IgG2a isotype control (Bio X Cell, BE0089, clone 2A3; RRID: AB_1107769) 6 hours after partial hepatectomy or CCl4 treatment. Depletion was confirmed by flow cytometric analysis of dissociated liver, blood, and bone marrow samples collected at the indicated time points.

### CCl4 and acetaminophen treatments

CCl4 (Sigma Aldrich, 289116) was diluted 1:10 in olive oil (Sigma Aldrich, O1514), and 10 μL/g body weight was injected intraperitoneally in 8- to 10-week-old mice^32^. Blood, bone marrow, and liver tissues were harvested at 24, 48, and 72 hours after the CCl4 injection. Acetaminophen (APAP, Sigma Aldrich, A7085) was given as a single dose in 10% DMSO in phosphate-buffered saline (PBS) and injected intraperitoneally at 300 mg/kg body weight^33^.

### Adenovirus delivery

For LIFR gain-of-function studies, 8-week-old C57BL/6J mice were injected with adenovirus (100 μL of the Adeno-TBG control virus or the Adeno-TBG-LIFR virus^29^, 5 × 10^8^ PFU per mouse, in saline) through the tail vein. At 5 or 10 days after adenovirus injection, 2/3 partial hepatectomy or CCl4 treatment was performed as described above.

### Primary hepatocyte isolation

The isolation of mouse hepatocytes was described previously^83^. Briefly, primary hepatocytes were isolated by perfusion of the mouse liver with perfusion buffer (5 mM EGTA in HBSS), followed by digestion with buffer containing 0.03-gram collagenase Ⅰ (Worthington Biochemical, LS004196) in 50 mL HBSS with 5 mM CaCl2. After digestion, the liver cells were filtered through a 70 μm cell strainer, followed by centrifugation at 50 × *g* for 3 min. Live hepatocytes were isolated by centrifugation in Percoll solution (0.48 mL 10× PBS + 4.32 mL Percoll + 5 mL DMEM) for 10 min at 800 rpm and then seeded on collagen (Sigma-Aldrich, 08-115)-coated plates in high-glucose DMEM containing 10% fetal bovine serum and 1% penicillin/streptomycin for 3 hours, followed by treatment with 100 ng/mL of HGF (Sino Biological, 50038-MNAH-20) or 20 ng/mL of EGF (Sino Biological, 50482-M01H-100) for 48 hours^84^. For *in vitro* adenoviral infection, hepatocytes isolated from C57BL/6J mice were incubated with control adenovirus or LIFR-expressing adenovirus for 24 hours before HGF or EGF treatment.

For measurements of Cxcl1 and cholesterol in the conditioned medium of primary hepatocytes, 5 × 10^5^ hepatocytes isolated from *Lifr*^flox/flox^ or *Lifr*^fl/fl^;Alb-Cre mice were cultured in each well of 6-well plates with 1.5 mL of high-glucose DMEM containing 10% fetal bovine serum and 1% penicillin/streptomycin for 48 hours.

### Quantitative reverse-transcription PCR

Total RNA was extracted by using TRIzol Reagent (Life Technologies, 15596018) or the PuroLink RNA Mini Kit (Invitrogen, 12183018A), and then reversed-transcribed with the iScript Reverse Transcription Supermix (Bio-Rad, 1708841). Quantitative PCR was performed with the iTaq Universal SYBR Green Supermix (Bio-Rad, 1725124) on a CFX96 real-time PCR machine (Bio-Rad). The mRNA level was calculated by using the ΔCt method and normalized to the internal control *Gapdh*. Sequences for qPCR primers are listed in **Supplementary Table 2**.

### Immunoblotting

Cultured cells or homogenized mouse tissues were lysed in RIPA lysis buffer (Sigma-Aldrich, 20- 188) containing protease inhibitors (GenDEPOT, P3100-001) and phosphatase inhibitors (GenDEPOT, P3200-001). Proteins were resolved on 4%-12% (GenScript, M00654) or 4%-20% (GenScript, M00657) precast gradient gels and transferred to a nitrocellulose membrane using the Trans-Blot Turbo Transfer System (Bio-Rad, 1704150). Membranes were blocked with 10% non-fat milk in Tris-buffered saline with 0.1% Tween 2(TBST) and incubated with the primary antibody at 4°C overnight, followed by incubation with the secondary antibody conjugated with horseradish peroxidase. The bands were visualized with an enhanced chemiluminescence substrate (ThermoFisher Scientific, 34578). Primary antibodies used are as follows: antibodies against Lifr (1:1,000, Proteintech, 22779-1-AP, RRID: AB_2879165), Gapdh (1:2,000, Proteintech, 60004-1- IG, RRID: AB_2107436), cyclin D1 (1:1,000, Cell Signaling Technology, 55506S, RRID: AB_2827374), cyclin A2 (1:1000, Proteintech, 18202-1-AP, RRID: AB_10597084), phospho- Stat3 (Tyr705) (1:1,000, Cell Signaling Technology, 9145S, RRID: AB_2491009), and Stat3 (1:1,000, Cell Signaling Technology, 30835S, RRID: AB_2798995). Immunoblotting images were obtained with the ChemiDoc Touch Imaging System (Bio-Rad) and Image Lab Touch software (Bio-Rad, version 2.3.0.07).

### Flow cytometry

For isolation and enrichment of liver-infiltrating leukocytes, liver tissues were digested and dissociated on the gentleMACS Dissociator with the Mouse Liver Dissociation Kit (Miltenyi Biotec, 130105807). After digestion and dissociation, the liver cells were filtered through a 100- μm cell strainer. Leukocytes were enriched by centrifugation with 40% Percoll plus (Millipore Sigma, Cytiva 17-5445-02). Blood samples were collected from the heart with a 1-mL syringe. Bone marrow samples were harvested from the tibia as described previously^85^. Cells were processed for surface marker staining as described previously^42^. Briefly, cells were depleted of red blood cells by using RBC Lysis Buffer (BioLegend, 420301), and 1 × 10^6^ cells per sample were used for staining. For dead cell staining, cells were incubated with Zombie Aqua Fixable Viability Kit (BioLegend, 423114) for 15 min. Cells were Fc-blocked with an anti-CD16/CD32 antibody (BioLegend, 101320) for 10 min and then incubated with a surface antibody mix for 30 min. After staining, cells were analyzed on an Invitrogen Attune NxT Acoustic Focusing Cytometer and analyzed by FlowJo software. We gated neutrophils by using ZombieDye^−^CD45^+^CD11b^+^Ly6G^+^. Antibodies used for flow cytometry staining are listed in **Supplementary Table 3**.

### CyTOF

CyTOF experiments were performed as described previously^86^. Briefly, liver tissues were cut into small pieces and dissociated on the gentleMACS Dissociator with the Mouse Liver Dissociation Kit (Miltenyi Biotec, 130105807). Leukocytes were enriched by centrifugation with 40% Percoll, and 1 × 10^6^ cells per sample were used for staining. For dead cell staining, cells were incubated with cisplatin (2.5 μM, Fisher Scientific, NC0637801) for 1 min. Cells were Fc-blocked with an anti-CD16/CD32 antibody (BioLegend, 101320) for 10 min and then incubated with a CyTOF surface antibody mix for 30-60 min. For intracellular staining, cells were incubated with fixation buffer (BioLegend, 420801) for 30 min and incubated in permeabilization buffer (BioLegend, 421002) at 4℃ overnight, followed by incubation with the CyTOF intracellular antibody mix for 30-60 min. For singlet discrimination, cells were washed and incubated with Cell-ID Intercalator-Ir (Fluidigm, 201192A) at 4 °C overnight. The samples were submitted to the Flow Cytometry and Cellular Imaging Core Facility at MD Anderson Cancer Center and run on CyTOF Instrumentation (DVS Science). CyTOF data were analyzed by Cytobank software (https://www.beckman.com/flow-cytometry/software/cytobank-premium). Polymorphonuclear neutrophils were identified as CD45^+^CD11b^+^Ly6G^+^. Kupffer cells were identified as CD45^+^CD11b^lo^F4/80^hi^. Monocyte-derived macrophages were identified as CD45^+^CD11b^hi^F4/80^lo^. CD8^+^ T cells were identified as CD45^+^CD3^+^CD8^+^. CD4^+^ T cells were identified as CD45^+^CD3^+^CD4^+^. B cells were identified as CD45^+^CD19^+^. Natural killer cells were identified as CD45^+^CD3^-^NK1.1^+^. Dendritic cells were identified as CD45^+^CD11c^+^MHC-II^+^. Eosinophils were identified as CD45^+^CD11b^+^Ly6G^-^Siglec-F^+^. Antibodies used for CyTOF are listed in **Supplementary Table 4**.

### Immunohistochemical and immunofluorescence staining

For these experiments, mice were euthanized and the livers were fixed in 10% neutral-buffered formalin (ThermoFisher Scientific) overnight, washed with PBS, transferred to 70% ethanol, embedded in paraffin, sectioned (5 μm thick), and stained with hematoxylin and eosin. Slides were deparaffinized in xylene and degraded alcohols. Heat-induced epitope retrieval was performed with a 2100-Retriever. Slides were rinsed with PBS, and a hydrophobic barrier was created around the tissue using a hydrophobic barrier pen (Vector Laboratories, H-4000-2). For immunohistochemical (IHC) staining, slides were placed in an incubating chamber with blocking solution (Vector Laboratories, SP-6000) for 10 min and rinsed with PBS, followed by incubation with 20% horse serum (Vector Laboratories, PK-7200) for 20 min. Next, slides were incubated with the primary antibody at 4 °C and rinsed with PBS, followed by incubation with a biotinylated universal secondary antibody (Vector Laboratories, PK-7200) for 30 min. After being washed again with PBS, slides were incubated with the avidin-biotin detection complex (ABC; Vector Laboratories, SK-4100) for 30 min and were then developed with 3,3′-diaminobenzidine (DAB) solution (Vector Laboratories, SK-4100). Counterstaining was done with Hematoxylin QS (Vector Laboratories, H-3404). The finished slides were scanned by the Leica Biosystems APERIO CS2. For immunofluorescence (IF) staining, slides were placed in an incubating chamber with blocking solution (3% BSA, 3% donkey serum, and 0.01% Triton in PBS) for 1 hour, followed by incubation with primary antibodies at 4 °C overnight. After being washed with PBS, slides were incubated with a secondary antibody for 1 hour. Slides were mounted with mounting medium with 4′,6- diamidino-2-phenylindole (DAPI) (Vector Laboratories, H-1200) and sealed with coverslips. Primary antibodies used for IHC and IF are as follows: antibodies against Ki67 (1:1,000, Bethyl Laboratories, IHC-00375; RRID: AB_1547959) and Ly6G (1:200, BioLegend, 127601; RRID:

AB_1089179). Secondary antibodies used for IF are as follows: donkey anti-rat IgG (H+L) highly cross-adsorbed secondary antibody (Alexa Fluor 488, 1:500, Thermo Fisher Scientific, A21208; RRID: AB_2535794), and donkey anti-rabbit IgG (H+L) highly cross-adsorbed secondary antibody (Alexa Fluor Plus 555, 1:500, Thermo Fisher Scientific, A32794; RRID: AB_2762834). The finished slides were scanned with a Vectra Polaris Automated Quantitative Pathology Imaging System. The numbers of positive cells were quantified with Image J software.

### Liver function analysis and TUNEL staining

Blood was collected from the mouse heart and centrifuged at 7,000 rpm at 4 ℃ for 15 min. Liver function was assessed by measuring the concentration of alanine aminotransferase (ALT) and aspartate aminotransferase (AST) in the serum by using an ALT Activity Assay Kit (Sigma-Aldrich, MAK052) and an AST Activity Assay Kit (Sigma-Aldrich, MAK055) according to the manufacturer’s protocols. Terminal deoxynucleotidyltransferase-mediated deoxyuridine triphosphate nick-end labeling (TUNEL) staining of paraffin-embedded tissue sections was done with the DeadEnd Fluorometric TUNEL System (G3250, Promega). TUNEL-positive areas from three fields (magnification×10) per animal were quantified with ImageJ software.

### Neutrophil isolation and staining

Mouse neutrophilic HGF was analyzed in blood samples or liver-infiltrating leukocytes of mice at 72 hours after partial hepatectomy, as described previously^48^. Briefly, samples were fixed by using fixation buffer (BioLegend, 420801) for 20 min, followed by Fc receptor blockade for 10 min and then incubation in permeabilization buffer (BioLegend, 421002) for 15 min. Cells were then stained with anti-HGF (R&D Systems, AF2207; RRID: AB_2118605) or IgG isotype control in permeabilization buffer for 30 min, followed by incubation with the donkey anti-goat IgG (H+L) highly cross-adsorbed secondary antibody, Alexa Fluor Plus 488 (Thermo Fisher Scientific, A32814TR; RRID: AB_2866497) for 30 min. Subsequently, surface markers were stained at 4°C for 30 min by using anti-CD45, anti-CD11b, and anti-Ly6G antibodies. Samples were analyzed on an Invitrogen Attune NxT Acoustic Focusing Cytometer with FlowJo software.

For measurement of HGF in the conditioned medium of purified neutrophils, neutrophils were isolated from the peripheral blood of mice as described previously^43^. Briefly, whole blood was collected via cardiac puncture (0.5 mL per animal) and suspended in HBSS (2 mL per animal) with 15 mM EDTA. After centrifugation (400 × *g*, 4 ℃, 10 min), cells were resuspended in 2 mL HBSS with 2 mM EDTA and then centrifuged (1,500 × *g*, room temperature, 30 min) in a three-layer Percoll gradient (78%, 69%, and 52%) without braking. Neutrophils enriched in the interface of 69% and 78% layers were confirmed to be of >90% purity by flow cytometry. Liver-infiltrating neutrophils were enriched at least twice by using the Neutrophil Isolation Kit (Miltenyi Biotec, 130097658). 1 × 10^5^ neutrophils were cultured in 100 µL RPMI 1640 medium containing 0.2% BSA for 6 hours.

Human polymorphonuclear neutrophils were isolated from healthy human blood obtained from the blood donation center of MD Anderson Cancer Center as described previously^87^. Briefly, blood was diluted at 1:1 with room-temperature PBS into a flask and mixed gently. To each Falcon tube pre-loaded with 15 mL of Ficoll solution (Cytiva, 17-1440-02), 35 mL of diluted blood was added slowly and then centrifuged at 800 × *g* at room temperature for 20 min. After centrifugation, the following layers were visible from the top: (1) plasma; (2) lymphocytes and monocytes (peripheral blood mononuclear cells); (3) Ficoll; (4) erythrocytes and granulocytes. The upper layer was removed by aspiration, and erythrocytes were lysed with ice-cold RBC Lysis Buffer (BioLegend, 420301). The purity of isolated polymorphonuclear neutrophils was greater than 95%, as gauged by flow cytometry and Giemsa (Sigma-Aldrich, GS500) staining (**Fig. 6k**). Purified polymorphonuclear neutrophils were added to 96-well plates and treated with cholesterol (Sigma-Aldrich, C4951) at the indicated concentrations for 4 hours in 100 µL of RPMI 1640 medium containing 0.2% BSA with Brefeldin A Solution (BioLegend, 420601). After treatment, samples were fixed with fixation buffer (BioLegend, 420801) for 20 min, followed by Fc receptor blockade for 10 min and then incubation in permeabilization buffer (BioLegend, 421002) for 15 min. Cells were then stained with anti-HGF (R&D Systems, MAB294; RRID: AB_2279754) or IgG isotype control antibody in permeabilization buffer for 30 min, followed by incubation with the goat anti-mouse IgG (H+L) cross-adsorbed secondary antibody, Alexa Fluor Plus 488 (Thermo Fisher Scientific, A-11001; RRID: AB_2534069) for 30 min. Samples were analyzed on an Invitrogen Attune NxT Acoustic Focusing Cytometer with FlowJo software.

### ELISA and cholesterol measurement

Enzyme-linked immunosorbent assay (ELISA) kits were used to measure the levels of mouse HGF (Sigma-Aldrich, RAB0214), mouse EGF (Sigma-Aldrich, RAB0150), mouse IL-6 (Sigma-Aldrich, RAB0308), mouse TNF-α (Cayman, Item No. 500850), mouse TGF-β (Proteintech, KE10005), and mouse Cxcl1 (Sigma-Aldrich, RAB0117) according to the manufacturer’s protocols. Free and total forms of cholesterol were measured by using the Cholesterol Quantitation Kit (Sigma-Aldrich, MAK043) according to the manufacturer’s protocols.

### Untargeted lipidomics analysis using LC-MS/MS

The same volumes of plasma samples (50 µL per mouse) collected from *Lifr*^fl/fl^ and *Lifr*^fl/fl^;Alb-Cre mice 72 hours after 2/3 partial hepatectomy were used for lipidomic analysis by the Metabolomics Core of MD Anderson Cancer Center as follows. To each plasma sample, 200 µL of extraction solution containing 2% Avanti SPLASH LIPIDOMIX Mass Spec Standard and 1% 10 mM butylated hydroxytoluene in ethanol was added and the tubes were vortexed for 10 min. The tubes were placed on ice for 10 min and centrifuged at 13,300 rpm at 4 ℃ for 10 min. The supernatant was transferred to a glass autosampler vial and the injection volume was 10 µL. Mobile phase A (MPA) was 40:60 for acetonitrile:water with 0.1 % formic acid and 10 mM ammonium formate. Mobile phase B (MPB) was 90:9:1 for isopropanol:acetonitrile:water with 0.1 % formic acid and 10 mM ammonium formate. The chromatographic method included a Thermo Fisher Scientific Accucore C30 column (2.6 µm, 150 × 2.1 mm) maintained at 40 °C, an autosampler tray chilled at 8 °C, a mobile phase flow rate of 0.200 mL/min, and a gradient elution program as follows: 0-3 min, 30% MPB; 3-13 min, 30-43% MPB; 13.1-33 min, 50-70% MPB; 48-55 min, 99% MPB; 55.1-60 min, 30% MPB.

A Thermo Fisher Scientific Orbitrap Fusion Lumos Tribrid mass spectrometer with a heated electrospray ionization source was operated in data-dependent acquisition mode, in both positive and negative ionization modes, with scan ranges of 150-827 and 825-1500 *m/z*. An Orbitrap resolution of 120,000 full width at half maximum (FWHM) was used for MS1 acquisition, and spray voltages of 3,600 V and -2900 V were used for positive and negative ionization modes, respectively. Vaporizer and ion transfer tube temperatures were set at 275 °C and 300 °C, respectively. The sheath, auxiliary, and sweep gas pressures were 35, 10, and 0 (arbitrary units), respectively. For MS^2^ and MS^3^ fragmentation, a hybridized HCD/CID approach was used. Each sample was analyzed using four injections making use of the two aforementioned scan ranges, in both ionization modes. Lipid data were processed and annotated by using Thermo Scientific LipidSearch software (version 5.0), and data were analyzed using R scripts written in-house. The Z-score was calculated as follows: the peak area (raw relative abundance) of each metabolite is normalized by subtracting the mean value of the metabolite across all samples, followed by division by the metabolite’s standard deviation.

### Statistical analysis

The statistical analysis used for each plot is described in the figure legends. Unless otherwise noted, data are presented as mean ± s.e.m, and Student’s *t*-test (two-tailed) was used to compare two groups of independent samples. The data analyzed by the *t*-test were normally distributed; we used an F-test to compare variances, and the variances were not significantly different. Therefore, when using a *t*-test, we assumed equal variance, and no data points were excluded from the analysis. *P* < 0.05 was considered statistically significant. Statistical analyses were done with GraphPad Prism (GraphPad Software, version 8).

## Acknowledgements

We thank MD Anderson’s Flow Cytometry and Cellular Imaging Core, Metabolomics Core, Functional Genomics Core, Cytogenetics and Cell Authentication Core, and Advanced Technology Genome Core for technical assistance. We are grateful to all members of the Ma Lab for the discussion and to Christine F. Wogan (MD Anderson’s Division of Radiation Oncology) for the critical reading of the manuscript. L.M. is supported by US National Institutes of Health (NIH) grants R01CA166051 and R01CA269140, a Cancer Prevention and Research Institute of Texas (CPRIT) grant RP190029, an American Cancer Society grant (award number: DBG-22- 161-01-MM), and the Nylene Eckles Distinguished Professorship of MD Anderson Cancer Center. H.Z. is supported by the NIH (R01AA028791, R01DK125396), CPRIT (RP220614), the Emerging Leader Award from the Mark Foundation For Cancer Research (award number: 21-003- ELA), and the Kern Wildenthal, M.D., Ph.D. Distinguished Professorship of University of Texas Southwestern Medical Center. The core facilities are supported by MD Anderson’s Cancer Center Support Grant P30CA016672 from the NIH.

## Author contributions

Y.D., Y.S., and L.M. conceived and designed the study. Y.D. and Z.Z. performed most experiments and data analyses. M.S., Y.Z., H.T., C.M., and J.Z. performed some experiments. C.L. and F.Y. provided technical assistance and consultation. Y.S. generated some reagents and contributed protocols. M.A.C. and H.Z. reviewed the data and provided significant intellectual input. Y.D. and L.M. wrote the manuscript with input from all other authors. L.M. provided scientific direction, established collaborations, and allocated funding for this study.

## Additional information

Extended Data: 5 figures. Supplementary information: 4 tables.

Correspondence and requests for materials should be addressed to L.M.

## Extended Data Figure Legends

**Extended Data Figure 1.**
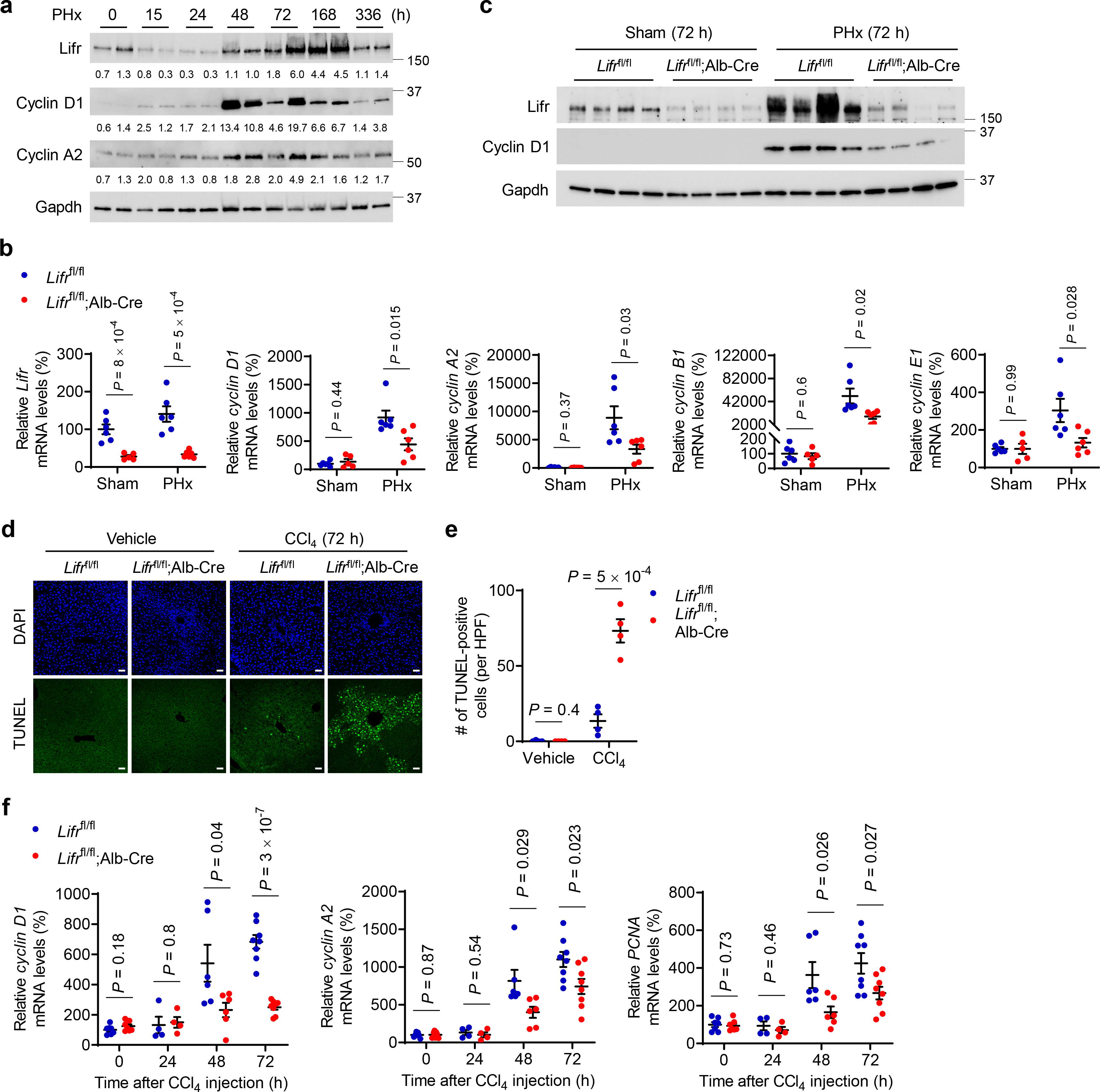
Loss of Lifr in hepatocytes impairs liver injury repair and injury-induced upregulation of proliferative genes. **a.** Immunoblotting of Lifr, cyclin D1, cyclin A2, and Gapdh in mouse livers at different time points after 2/3 partial hepatectomy (PHx). **b.** qPCR of mRNA of Lifr, cyclin D1, cyclin A2, cyclin B1, and cyclin E1 in the livers of *Lifr*^fl/fl^ and *Lifr*^fl/fl^;Alb-Cre mice at 72 hours after PHx. *n* = 6, 5, 6, and 6 mice. **c.** Immunoblotting of Lifr, cyclin D1, and Gapdh in the livers of *Lifr*^fl/fl^ and *Lifr*^fl/fl^;Alb-Cre mice at 72 hours after PHx. **d**, **e.** Representative TUNEL staining (**d**) and the number of TUNEL-positive hepatocytes per high-power field (HPF; **e**) at 72 hours after CCl4 treatment. Scale bars, 50 μm. *n* = 4 mice per group. **f.** qPCR of mRNA of cyclin D1, cyclin A2, and PCNA in the livers of *Lifr*^fl/fl^ and *Lifr*^fl/fl^;Alb-Cre mice at the indicated times after CCl4 treatment. *n* = 7, 7, 4, 4, 6, 6, 8, and 8 mice. Statistical significance in **b**, **e**, and **f** was determined by a two-tailed unpaired *t*-test. Error bars are s.e.m.

**Extended Data Figure 2.**
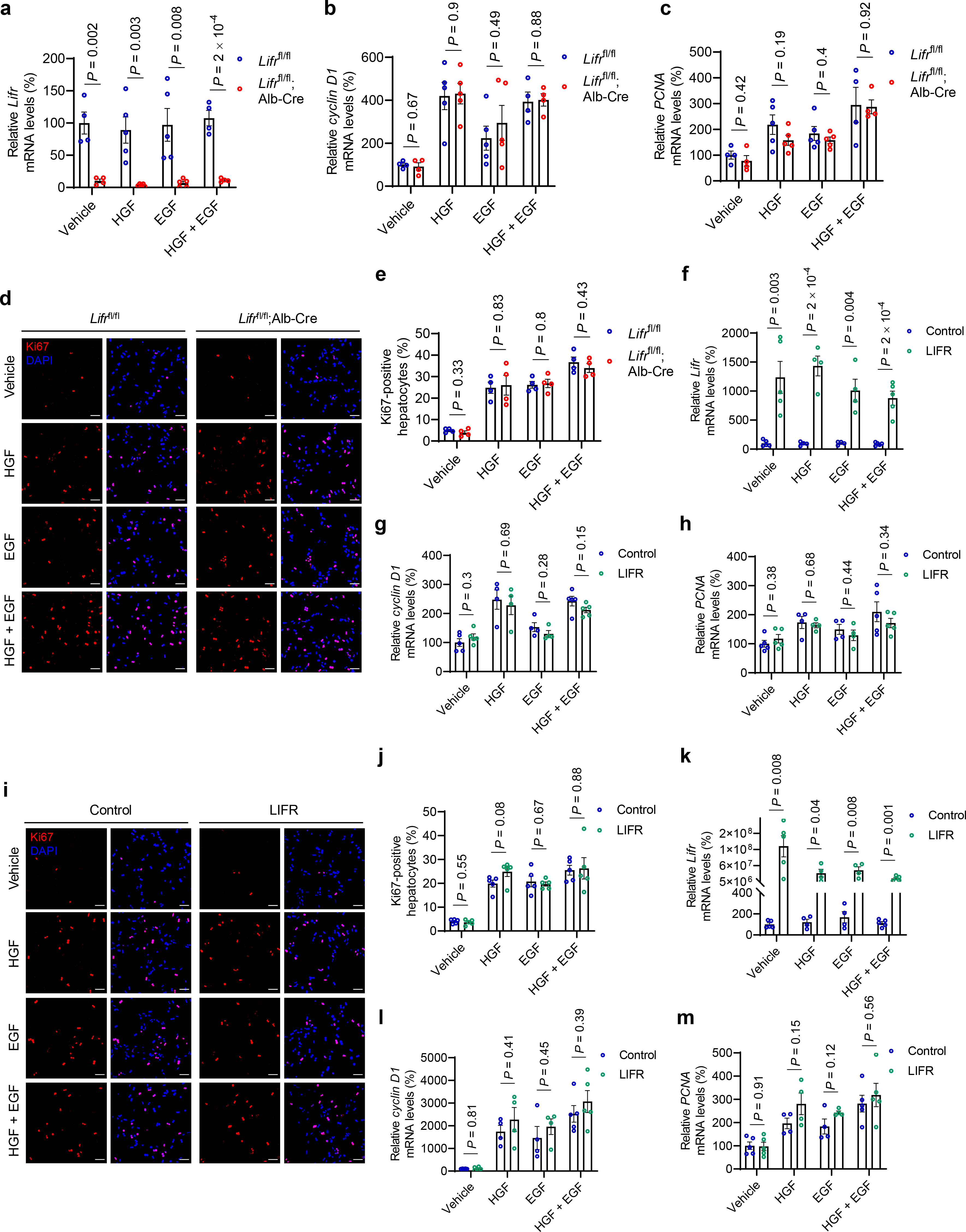
LIFR deficiency or overexpression does not affect hepatocyte proliferation *ex vivo* or *in vitro*. **a-e.** Primary hepatocytes isolated from *Lifr*^fl/fl^ and *Lifr*^fl/fl^;Alb-Cre mice were cultured for 3 hours, followed by treatment with 100 ng/mL of HGF and/or 20 ng/mL of EGF for 48 hours. **a-c.** qPCR of mRNA of Lifr (**a**), cyclin D1 (**b**), and PCNA (**c**) in HGF- and/or EGF-treated hepatocytes. *n* = 4, 4, 5, 5, 5, 5, 4, and 4 biological replicates. **d**, **e.** Immunofluorescence staining of Ki67 (**d**; overlay with DAPI staining) and percentage of Ki67-positive cells (**e**) in HGF- and/or EGF-treated hepatocytes. *n* = 4 biological replicates per group. **f**-**j.** Primary hepatocytes isolated from C57BL/6J mice 10 days after injection with control adenovirus or LIFR-expressing adenovirus were cultured for 3 hours, followed by treatment with 100 ng/mL of HGF and/or 20 ng/mL of EGF for 48 hours. **f-h.** qPCR of mRNA of Lifr (**f**), cyclin D1 (**g**), and PCNA (**h**) in HGF- and/or EGF-treated hepatocytes. *n* = 5, 5, 4, 4, 4, 4, 5, and 5 biological replicates. **i**, **j.** Immunofluorescence staining of Ki67 (**i**; overlay with DAPI staining) and percentage of Ki67- positive cells (**j**) in HGF- and/or EGF-treated hepatocytes. *n* = 5 biological replicates per group. **k**-**m.** qPCR of mRNA of Lifr (**k**), cyclin D1 (**l**), and PCNA (**m**) in HGF- and/or EGF-treated hepatocytes isolated from C57BL/6J mice. The cells were infected with control adenovirus or LIFR-expressing adenovirus for 24 hours before HGF and/or EGF treatment. *n* = 5, 5, 4, 4, 4, 4, 5, and 5 biological replicates. Statistical significance in **a**-**c**, **e**-**h**, and **j-m** was determined by a two-tailed unpaired *t*-test. Error bars are s.e.m.

**Extended Data Figure 3.**
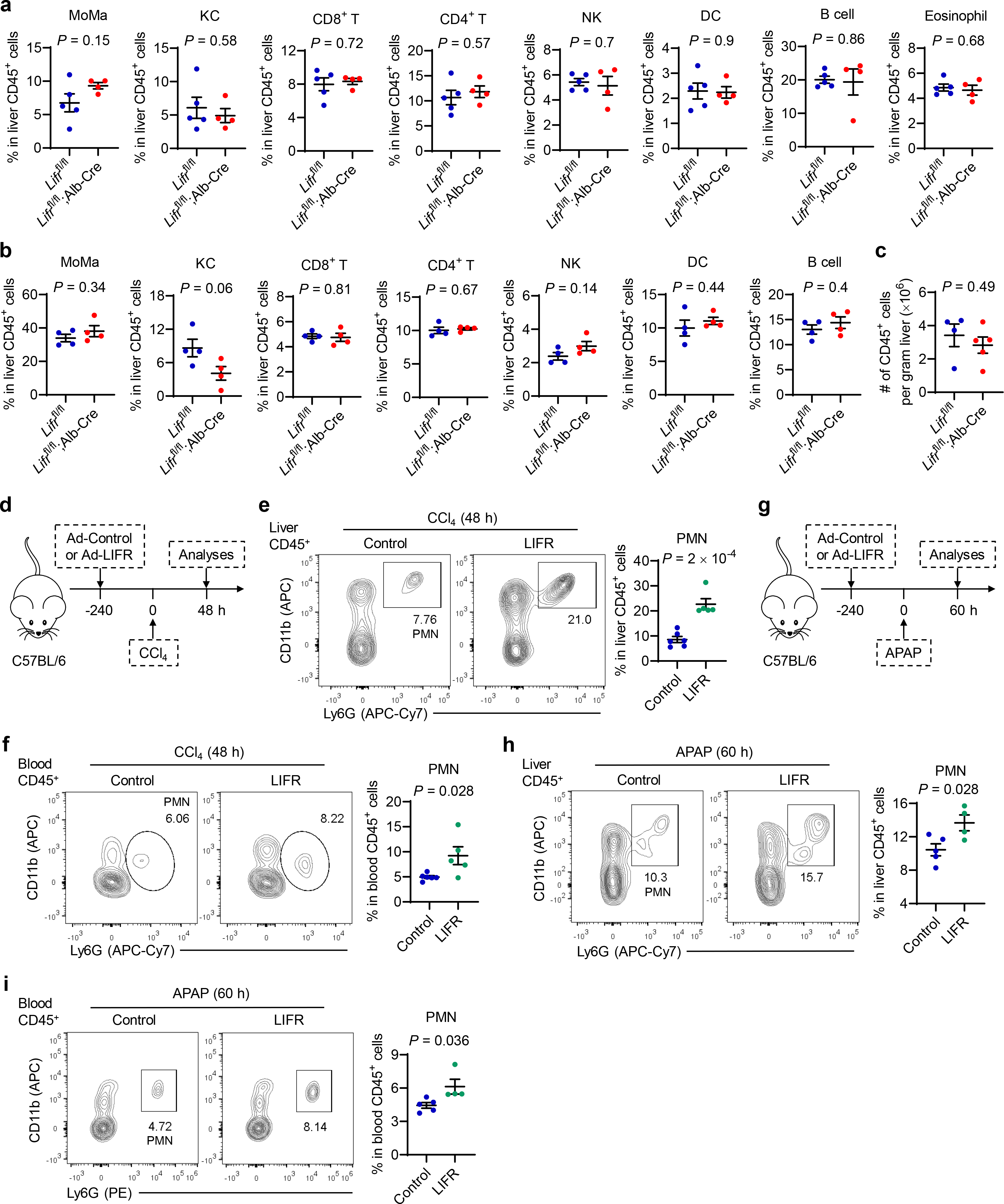
Effects of LIFR on neutrophil recruitment in CCl4- and acetaminophen-induced liver injury models. **a, b.** Quantification of liver-infiltrating immune cell populations in *Lifr*^fl/fl^ and *Lifr*^fl/fl^;Alb-Cre mice 72 hours after PHx (**a**; *n* = 5 and 4 mice) or CCl4 treatment (**b**; *n* = 4 mice per group). NK: natural killer cells. KC: Kupffer cells. MoMa: monocyte-derived macrophages. PMN: polymorphonuclear neutrophils. DC: dendritic cells. **c.** Number of CD45^+^ cells per gram of liver in *Lifr*^fl/fl^ and *Lifr*^fl/fl^;Alb-Cre mice at 72 hours after PHx. *n* = 4 and 5 mice. **d**-**f.** C57BL/6J mice received control or LIFR-expressing adenovirus 10 days before CCl4 treatment. Analyses were done at 48 hours after CCl4 treatment. **d.** Schematic of the experimental design. **e**, **f.** Representative flow cytometry plots and percentage of neutrophils in liver (**e**) and blood (**f**) CD45^+^ cells from mice at 48 hours after CCl4 treatment. *n* = 6 and 5 mice. **g**-**i.** C57BL/6J mice received control or LIFR-expressing adenovirus 10 days before acetaminophen (APAP) treatment. Analyses were done at 60 hours after APAP treatment. **g.** Schematic of the experimental design. **h, i.** Representative flow cytometry plots and percentage of neutrophils in liver (**h**) and blood (**i**) CD45^+^ cells from mice at 60 hours after APAP treatment. *n* = 5 and 4 mice. Statistical significance in **a-c**, **e**, **f**, **h**, and **i** was determined by a two-tailed unpaired *t*-test. Error bars are s.e.m.

**Extended Data Figure 4.**
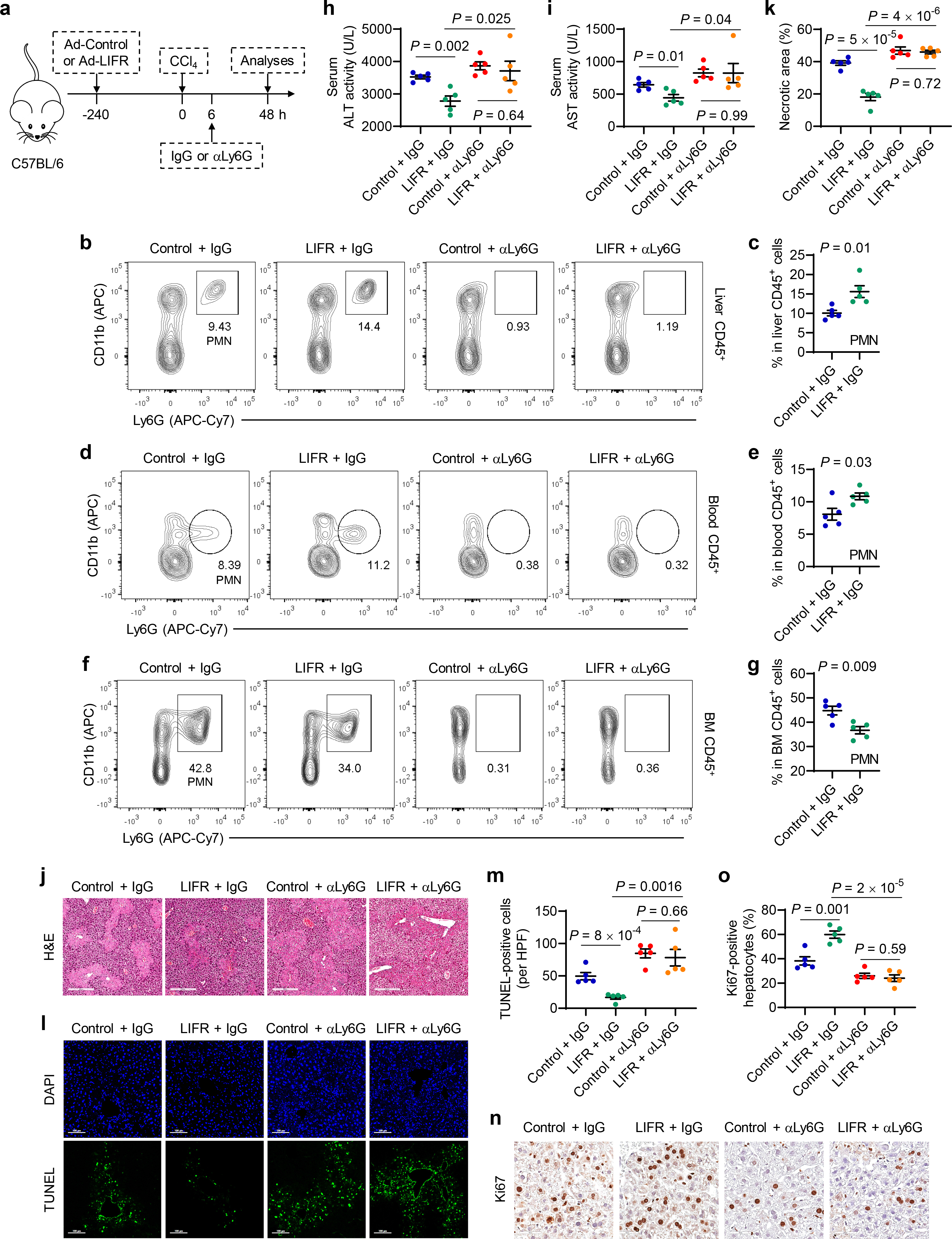
LIFR accelerates CCl4-induced liver injury repair and regeneration in a neutrophil-dependent manner. **a**-**n.** C57BL/6J mice received control or LIFR-expressing adenovirus 10 days before CCl4 treatment. 6 hours after CCl4 treatment, the mice were treated with control IgG or anti-Ly6G. Analyses were done at 48 hours after CCl4 treatment. **a.** Schematic of the experimental design. **b, c.** Representative flow cytometry plots (**b**) and percentage of neutrophils in liver CD45^+^ cells (**c**). *n* = 5 mice per group. **d, e.** Representative flow cytometry plots (**d**) and percentage of neutrophils in blood CD45^+^ cells (**e**). *n* = 5 mice per group. **f, g.** Representative flow cytometry plots (**f**) and percentage of neutrophils in bone marrow (BM) CD45^+^ cells (**g**). *n* = 5 mice per group. **h, i.** Serum ALT (**h**) and AST (**i**) levels in control and LIFR-expressing adenovirus-infected C57BL/6J mice injected with control IgG or anti-Ly6G after CCl4 treatment. *n* = 5 mice per group. **j, k.** Representative H&E staining (**j**) and percentage of necrotic areas (**k**). Scale bars, 300 μm. *n* = 5 mice per group. **l, m.** Representative TUNEL staining (**l**) and the number of TUNEL-positive hepatocytes per high-power field (HPF; **m**). Scale bars, 100 μm. *n* = 5 mice per group. **n, o.** Immunohistochemical staining of Ki67 (**n**) and percentage of Ki67-positive hepatocytes (**o**). Scale bars, 50 μm. *n* = 5 mice per group. Statistical significance in **c**, **e**, **g**, **h**, **i**, **k**, **m**, and **o** was determined by a two-tailed unpaired *t*-test. Error bars are s.e.m.

**Extended Data Figure 5.**
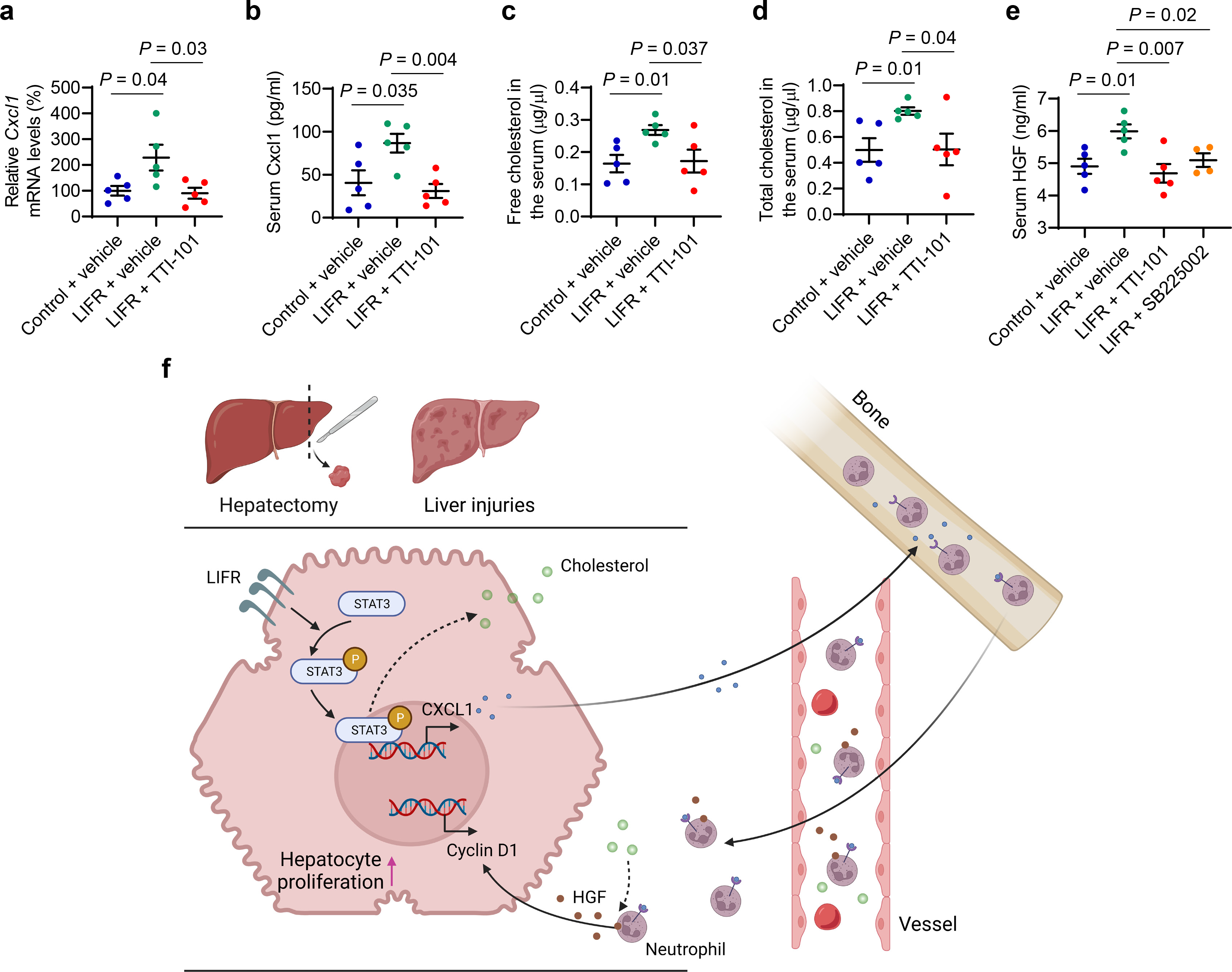
LIFR boosts the production of CXCL1, cholesterol, and HGF after liver injury in a STAT3-independent manner. **a, b.** qPCR of *Cxcl1* in the livers (**a**) and ELISA of secreted Cxcl1 in the serum (**b**) of C57BL/6J mice that received control or LIFR-expressing adenovirus 10 days before PHx and were treated with TTI-101 or SB225002 for 3 days. Analyses were done at 48 hours after PHx. *n* = 5 mice per group. **c, d.** Levels of free cholesterol (**c**) and total cholesterol (**d**) in the serum of the mice described in **a**. *n* = 5 mice per group. **e.** ELISA of HGF in the serum of the mice described in **a**. *n* = 5, 5, 5, and 4 mice per group. **f.** Model for LIFR-mediated regulation of neutrophil recruitment and liver regeneration. Statistical significance in **a-e** was determined by a two-tailed unpaired *t*-test. Error bars are s.e.m.

## Supplementary Tables

**Supplementary Table 1.** Normalized Z-scores of lipids in plasma samples collected from control and Lifr conditional knockout mice at 72 hours after partial hepatectomy This table is presented as a separate Excel file.

**Supplementary Table 2.**
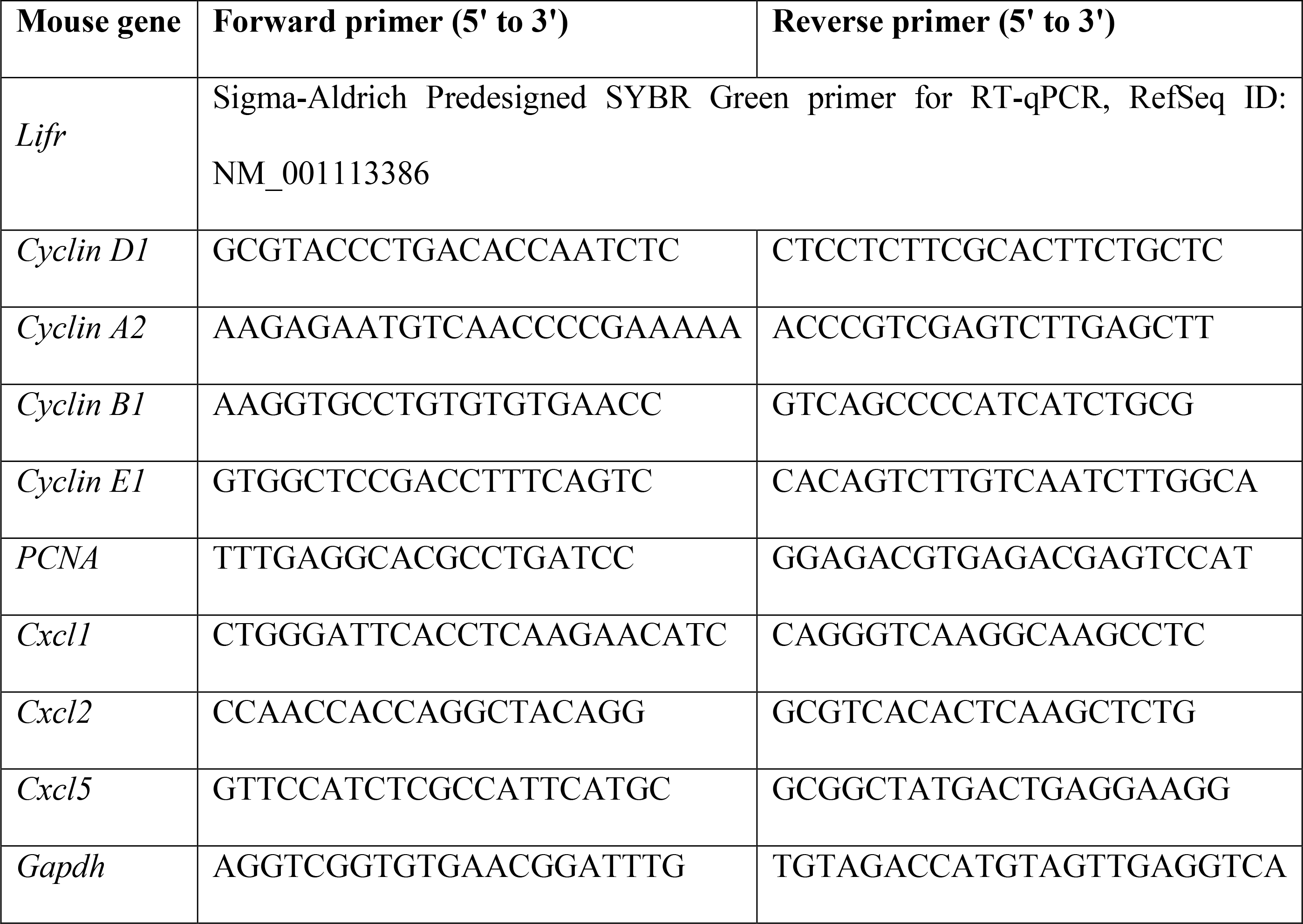
Primers used for qPCR

**Supplementary Table 3.**
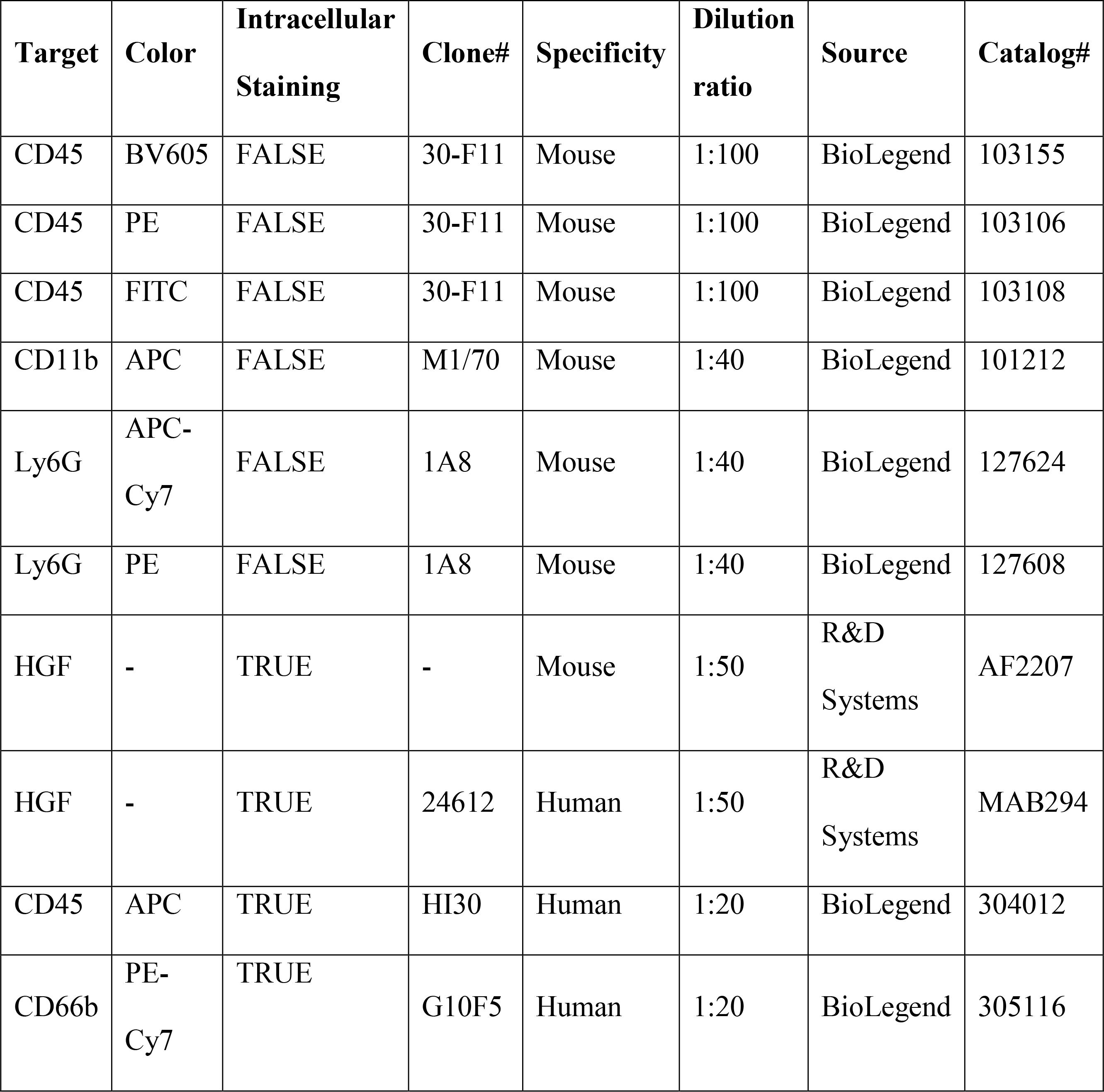
Antibodies used for flow cytometry

| **Target** | **Color** | **Intracellular Staining** | **Clone#** | **Specificity** | **Dilution ratio** | **Source** | **Catalog#** |
| --- | --- | --- | --- | --- | --- | --- | --- |
| CD45 | BV605 | FALSE | 30-F11 | Mouse | 1:100 | BioLegend | 103155 |
| CD45 | PE | FALSE | 30-F11 | Mouse | 1:100 | BioLegend | 103106 |
| CD45 | FITC | FALSE | 30-F11 | Mouse | 1:100 | BioLegend | 103108 |
| CD11b | APC | FALSE | M1/70 | Mouse | 1:40 | BioLegend | 101212 |
| Ly6G | APC-Cy7 | FALSE | 1A8 | Mouse | 1:40 | BioLegend | 127624 |
| Ly6G | PE | FALSE | 1A8 | Mouse | 1:40 | BioLegend | 127608 |
| HGF | - | TRUE | - | Mouse | 1:50 | R&D Systems | AF2207 |
| HGF | - | TRUE | 24612 | Human | 1:50 | R&D Systems | MAB294 |
| CD45 | APC | TRUE | HI30 | Human | 1:20 | BioLegend | 304012 |
| CD66b | PE-Cy7 | TRUE | G10F5 | Human | 1:20 | BioLegend | 305116 |

**Supplementary Table 4.** Antibodies used for CyTOF

| **Target** | **Label** | **Intracellular staining** | **Clone#** | **Dilution ratio** | **Source** | **Catalog#** |
| --- | --- | --- | --- | --- | --- | --- |
| CD45(Ms) | 89Y | FALSE | 30-F11 | 1:200 | DVS-Fluidigm | 3089005B |
| CD3, CD3e | 152Sm | FALSE | 145-2C11 | 1:250 | BioLegend | 100302 |
| CD4(Ms) | 145Nd | FALSE | RM4-5 | 1:400 | BioLegend | 100506 |
| CD8a | 115In | FALSE | 53-6.7 | 1:250 | BioLegend | 100702 |
| CD11b | 143Nd | FALSE | M1/70 | 1:400 | DVS-Fluidigm | 3143015B |
| Ly-6G | 146Nd | FALSE | 1A8 | 1:250 | BioLegend | 127637 |
| F4/80 | 174Yb | FALSE | BM8.1 | 1:333 | Tonbo | 70-4801-U100 |
| NK1.1 | 170Er | FALSE | PK136 | 1:200 | BioLegend | 108702 |
| CD19 | 149Sm | FALSE | 4D5 | 1:250 | BioLegend | 115502 |
| I-A/I-E, MHC-II | 209Bi | FALSE | M5/114.15.2 | 1:500 | DVS-Fluidigm | 3209006B |
| CD11c | 171Yb | FALSE | N418 | 1:200 | BioLegend | 117302 |
| Siglec-F | 172Yb | FALSE | E50-2440 | 1:250 | BD | 552125 |

